# Molecular evolution of luciferase diversified bioluminescent signals in sea fireflies

**DOI:** 10.1101/2020.01.23.917187

**Authors:** Nicholai M. Hensley, Emily A. Ellis, Nicole Y. Leung, John Coupart, Alexander Mikhailovsky, Daryl A. Taketa, Michael Tessler, David F. Gruber, Anthony W. De Tomaso, Yasuo Mitani, Trevor J. Rivers, Gretchen A. Gerrish, Elizabeth Torres, Todd H. Oakley

## Abstract

Understanding the genetic causes of evolutionary diversification is challenging because differences across species are complex, often involving many genes. However, cases where single or few genetic loci affect a feature that varies dramatically across a radiation of species would provide tractable opportunities to understand the genetics of diversification. Here, we show the diversification of bioluminescent signals in cypridinid ostracods (“sea fireflies”) to be strongly influenced by a single gene, cypridinid-luciferase. We find different evolutionary processes, including selection, drift, and constraint, each acted on c-luciferase at different times during evolutionary history and impacted different phenotypes, diversifying behavioral signals across species. In particular, some amino acid sites in c-luciferase evolved under episodic diversifying selection, and are associated significantly with phenotypic changes in both enzyme kinetics and color, which impact signals directly. We also find that multiple other amino acid positions in c-luciferase evolved neutrally or under purifying selection and may have impacted the variation of color of bioluminescent signals across genera. This work provides a rare glimpse into the genetic basis of diversification across many species, showing how multiple evolutionary processes may act at different times during a radiation of species to diversify phenotypes. These results indicate not only selection but also drift and constraint may be important evolutionary drivers of species diversification.

**Significance statement:** A hallmark of life is its astounding diversity. While we are beginning to understand the drivers of biodiversity, uncovering the genetic basis remains challenging. As such, how different molecular evolutionary processes act to diversify phenotypes is a major question in biology. Here we show a single gene to be important in a riotous diversity of fantastical behaviors - the bioluminescent signals of sea fireflies - allowing us to demonstrate multiple evolutionary forces including selection, drift, and constraint contributed to diversification. Our work highlights that not only selection but also neutral processes and constraint have each worked at different times to shape phenotypic diversity.

## Main

Why and how some groups of species diversify more than others are enduring questions in biology, with broad implications for the origin and maintenance of biodiversity. Particularly challenging is to understand the genetic underpinnings of diversification, because numerous genes typically underlie quantitative phenotypic differences that vary across many species (1–3). In addition, wide disparities across species also exist in particular traits like morphology and behaviors including courtship that often accompany rapid speciation (4–8). During rapid divergence of particular traits, we might expect few loci of large effect (e.g. effector genes *sensu (9)*) or with a simple genetic architecture (10) may be better poised to diversify phenotypes quickly (11, 12), but such genes are often difficult to identify. Rapidly diversifying phenotypes have therefore inspired genome-scale studies of species radiations, which show that positive and purifying selection are involved in species differences and sometimes can act on a limited number of loci to promote such variety (13, 14). When possible, another approach to the genetics of diversification is to identify cases where one or a few genes are linked to diverse phenotypes across many species. Visual pigment genes (opsins) are a prime example of a gene family that underlies an organismal phenotype (color sensitivity) with a shared genetic basis across species (15–17). While most genomic and single-gene studies highlight how selection and/or epistasis impact phenotypic diversification (14, 18), a role for neutral processes has been difficult to demonstrate (19).

Bioluminescent ostracods (family Cypridinidae) and their phenotypically disparate signals provide a particularly attractive system for integrative understanding of diversification (5, 20–23). Bioluminescent cypridinids globally use light as an anti-predator display, including Caribbean species that also use light for courtship. This Caribbean clade diversified into dozens of species, each with a courtship display that is conserved within, but variable between species (22, 24). Although both sexes use light for anti-predator displays by expelling light-producing chemicals mixed with mucus, as far as we know, only males of Caribbean species produce courtship signals. In contrast to the light cloud produced during anti-predator signals, courtship signals are comprised of delicate, discrete pulses of light formed as males rapidly swim (23, 25). Multiple species commonly live in geographical sympatry, yet produce signals above different microhabitats. These courtship signals also vary in other parameters, including display angle (with reference to the benthos), specific time of onset (26), and time and distance between light pulses (21). Pulses can vary in brightness, kinetics, and color (20, 23, 27). Once ostracods externally secrete the pulses in bioluminescent signals, the phenotype of a single pulse is dictated by biochemistry instead of behavior, and depends largely on well-understood chemical reactions between cypridinid luciferase (“c-luciferase”) and the substrate, luciferin. Because the substrate is shared within this ostracod family (27), biochemical differences in light production largely depend on differences in c-luciferases. Therefore, cypridinid ostracods provide a system whereby some critical aspects of signals may be separated from the complexities of behavior and connected directly to genetic changes in enzymes, allowing unique insights into the relationships between genotype, phenotype, and diversification.

The decay rate of light production is one target for connecting genotype, phenotype, and diversification of signals in cypridinid ostracods. Recent work shows decay rates vary considerably across species of luminous ostracods, and are correlated with the temporal duration of light in the individual pulses of courtship signals (20). Hensley et al. (2019) measured kinetics of light production by fitting an exponential decay parameter to each species’ bioluminescence. Cypridinid bioluminescence is a single-order biochemical reaction whose rate depends on the concentration of substrate (28). Although other components within the mucus, including varying levels of luciferin, alter the overall reaction dynamics, light production fades exponentially once substrate becomes limiting and is a reliable metric of c-luciferase function. We hypothesize that differences in sequences of c-luciferase enzymes influence differences in the light decay kinetics, which vary dramatically across species and affect the light duration in both anti-predation and courtship displays. The duration of anti-predator light could be under natural selection, and the duration of courtship pulses could be under sexual selection, perhaps used for species recognition, and/or choosing amongst potential conspecific mates.

In addition to kinetics of light decay, the color of cypridinid bioluminescence could be dictated by differences in c-luciferase proteins. However unlike kinetics, emission spectra (“colors”) are not well-characterized in many cypridinids. Previously published experiments hinted at variation in color of ostracod bioluminescence (27, 29), yet interspecific comparisons were impossible due to differences in methods and lack of replication (Table S1). Harvey (27) first noticed a difference in color between two species when he cross-reacted crude preparations from *V. hilgendorfii* and an unknown Jamaican species. He noted the luciferase preparation of *V. hilgendorfii* catalyzed a “bluish” light and the Jamaican luciferase a “yellowish” light, concluding that the protein (luciferase) dictated the color. The qualitative observations of Harvey nearly 100 years ago are consistent with the more recently published emission spectra of ostracods (29–32) that suggest some Caribbean species have higher λ_max_ values than other species. If so, λ_max_ may be another phenotype that varies across species and is dictated by c-luciferase. Like the rate of light decay, differences in color of bioluminescence could be functionally neutral, or could be influenced by natural selection, mediated through its appearance to would-be predators, and/or sexual selection, mediated through its appearance to would-be mates.

Because c-luciferases may be expressed *in vitro* to test biochemical functions (33) - including light production, decay kinetics, and color - this system fits squarely into the “functional synthesis” research paradigm (34), which combines phylogenetic analyses of sequences and manipulative biochemical experiments to allow mechanistic understanding of adaptation and evolutionary constraints. However, despite the potential value of bioluminescence for genotype-phenotype relationships (35, 36), previous studies that characterize luciferase genes of cypridinid bioluminescence exist for only two species (32, 37), which show cypridinid luciferases evolved independently of other luciferases, have a signal peptide leading to secretion outside cells, and possess two Von Willebrand Factor-D (VWD) domains (38). Here, we characterize new c-luciferase genes for phylogenetic sequence analyses in combination with functional characterizations of bioluminescence. In particular, we hypothesize that differences in biochemical kinetics, namely the rate of decay of light production (“decay”), and emission spectra of bioluminescence (“color”) can be linked to genetic variation in c-luciferase genes during the diversification of both anti-predator and courtship signals, to allow us to understand the molecular evolutionary processes that shaped

## Results

### Thirteen new putative luciferases

We identified 13 new putative luciferases from whole-body transcriptomes of cypridinid ostracods. These genes form part of a monophyletic family along with the only two previously published cypridinid luciferases (Fig. 1; Fig. S1). These putative luciferases bear multiple hallmarks of c-luciferase, including two Von Willebrand Factor-D (VWD) domains, a signal peptide, and conserved cysteine sites (32, 37, 38).

**Figure 1.**
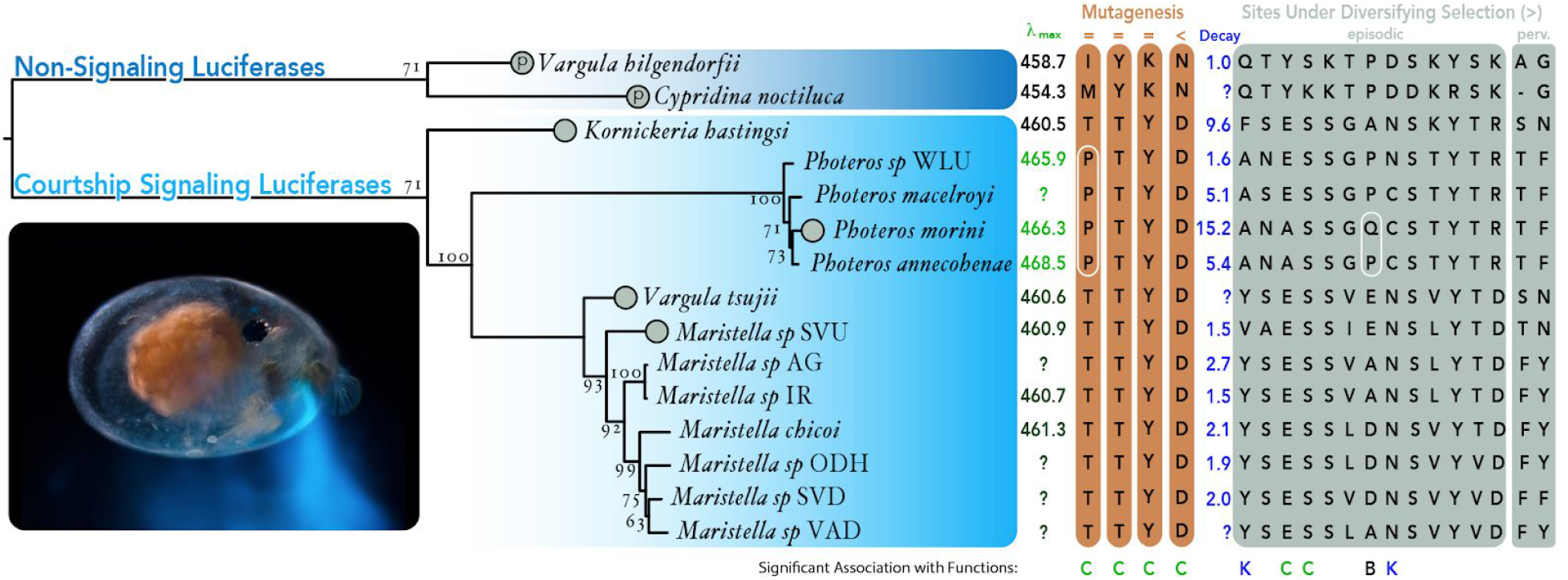
Maximum likelihood phylogeny of new and published c-luciferase genes using codon-aligned DNA and GTR model with fast bootstraps (percentages at nodes) in IQ-Tree. Circles at tips of phylogeny illustrate six genes expressed in vitro that show light catalysis. Two circles with “p” inside are previously published, the other four are explained herein (Fig 2). At right are two biochemical phenotypes (*λ_max_* and light decay constant), and interesting amino acid sites, including three that affect *λ_max_* in mutagenesis experiments (orange shading), thirteen under episodic diversifying selection using the MEME approach of hyphy and two under pervasive diversifying selection using the FEL approach of hyphy. Numbers of sites at bottom correspond to the unaligned *Cypridina noctiluca* luciferase sequence. We label the dn:ds ratio above the sites with “>” meaning diversifying/positive selection, “<” meaning purifying selection, and “=” meaning failure to reject null neutral model. White ellipses around sites are amino acid patterns discussed in the main text. We label sites significantly associated with Color (C), Kinetics (K), and Both color and kinetics (B). Table 1 has p-values for these sites. From left to right, site numbers are (*λ_max_* values), 38, 178, 375, 404, (Decay values), 19, 67, 74, 114, 132, 148, 160, 232, 256, 262, 289, 358, 445, 20, 180.

### Patterns of natural selection in c-luciferases

Using mixed effects maximum likelihood in HYPHY (39), we found ratios of synonymous to non-synonymous substitution rates in c-luciferases to indicate twelve sites consistent with episodic diversifying selection (Fig. 1), distributed throughout the gene. “Episodic” diversifying selection refers to sites with elevated dn:ds along a proportion of branches of the gene tree. According to Fixed Effects Likelihood (FEL) analysis (40), 250 c-luciferase sites have significantly low rates of dn:ds, consistent with purifying (negative) selection, and two sites with significantly high dn:ds, consistent with pervasive positive selection, or sites with elevated dn:ds for the entire gene tree.

### Luciferases from Exemplar Species are Functional

Using *in vitro* expression in mammalian and/or yeast cells (Fig 2), we tested the ability of putative c-luciferases to catalyze light reactions. We selected four exemplar species representing different genera of bioluminescent Cypridinidae. For each luciferase construct, light levels increased significantly after the addition of the substrate luciferin or compared to negative controls (Fig. 2; Table S2). Adding luciferin to biological media often produces light (41) that varies across biological replicates. We note this variation, yet also report statistically significant differences in light production after adding substrate (for yeast) or between c-luciferase secreting-cells and cells or media alone (for HEK293 cells and yeast) (Fig 2). This is consistent with the putative c-luciferases across multiple genera of Cypridinidae being functional c-luciferases.

**Figure 2.**
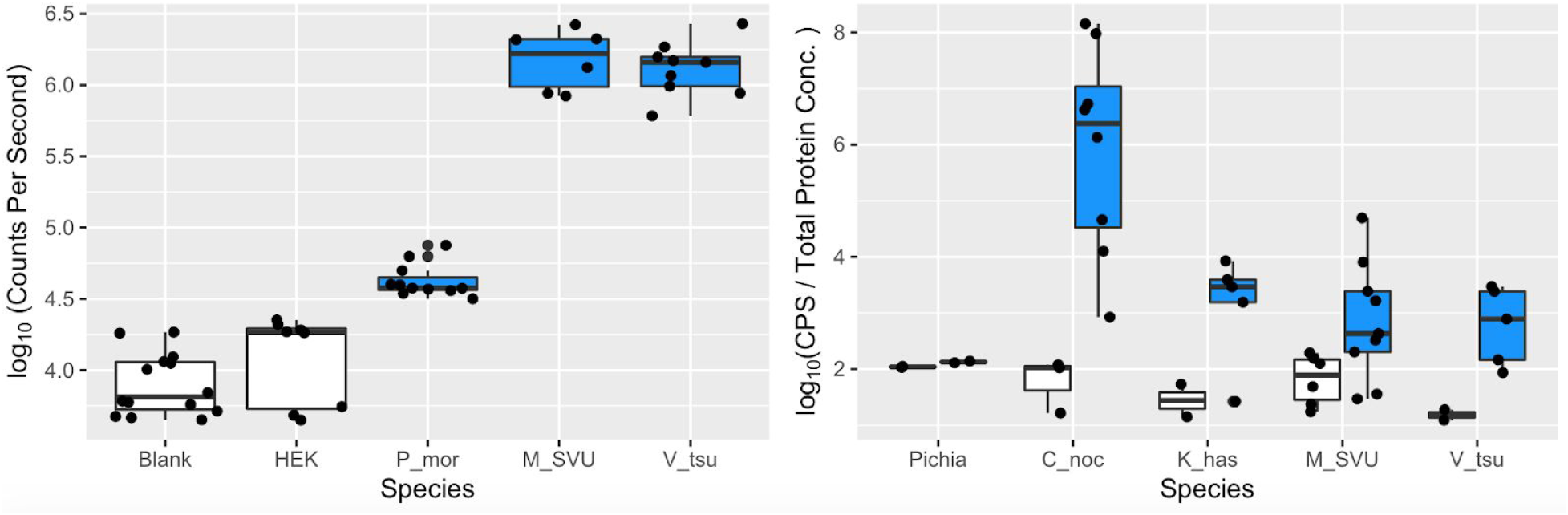
Exemplar luciferases are functional. **A**. Light produced after addition of luciferin to putative c-luciferase constructs from different species, measured in log Counts Per Second. Controls include blank cell culture media and HEK cells with no construct. Constructs are: (1) *Photeros morini* (P_mor), *Maristella sp SVU* (M_SVU), and *Vargula tsujii*. (V_tsu) **B**. Log counts per second normalized by total protein concentration for four c-luciferase constructs expressed in yeast *Pichia*. Measures are before the addition of the substrate luciferin (grey) and after (blue). Control is *Pichia* cells alone. Constructs are: (1) the host strain of *Pichia* without a luciferase as a negative control, (2) a sequence known from *Cypridina noctiluca* (C_noc) as a positive control, (3) a novel sequence from *Kornickeria hastingsi carriebowae* (K_has), (4) a novel sequence from an undescribed species from Belize, “SVU” (nominal genus *Maristella*), and (5) a novel sequence from the California sea firefly *Vargula tsujii*. Each datum is an average of three technical replicates.

### Emission spectra vary across species of bioluminescent Cypridinidae

By crushing whole specimens to elicit light production in front of a spectroradiometer (see Methods), we obtained new data on emission spectra from 20 species (Fig. 3; Tables S3, S4). The wavelength of maximum emission (λ_max_) varies from 458.7 ± 1.80 nm in *V. hilgendorfii* to 468.0 ± 1.80 nm in *Photeros sp. EGD*. Previously published data extend this range, with *Cypridina noctiluca* at 454.3 nm and *Photeros graminicola* at 471.1 nm. We found *λ_max_* from species of *Photeros* do not overlap in value with *λ_max_* of other species. The lowest *λ_max_* from any *Photeros* we measured is 465.6 ± 0.78 nm, whereas the highest of any non-Photeros species is 461.5 ± 2.70. Assuming *Photeros* is monophyletic, we infer an evolutionary increase in *λ_max_* along the branch leading to *Photeros*. Monophyly of *Photeros* is supported by published morphological (42) and molecular (20) phylogenetic analyses, by the phylogeny of c-luciferases we present here (Fig. 1), and by a diagnostic ratio of length:height of carapace (21, 43, 44), which we also used here to classify undescribed species into genera (Table S5). Full Width Half Maximum (FWHM) is a common parameter to describe the variation (width) of wavelengths in emission spectra. We found FWHM values to also vary between species of Cypridinidae (Fig. S2), ranging from 75.05 ± 0.91 to 85.44 ± 0.32, although we notice no apparent phylogenetic pattern.

**Figure 3.**
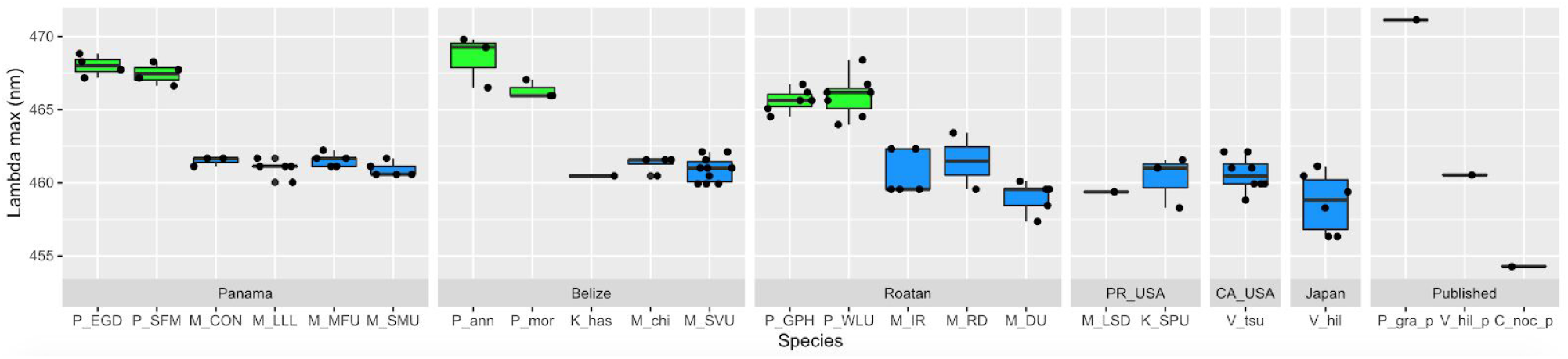
*Photeros* (green) have higher values of λ_max_ than other genera (blue). Wavelength value with highest emission (λ_max_ ; “Lambda max” on the y-axis) in nanometers (nm) from new emission spectra from 20 species and previously published spectra from 3 species.

### Genetic basis of phenotypic changes

By analyzing previously published mutagenesis studies along with new c-luciferase sequences and new color data, we identified amino acid sites that influence λ_max_. The site most strongly associated with λ_max_ is 178 (ANOVA, p=2.2 e-16; Table 1). This effect is clear from published mutations alone: wild-type luciferase from *Cypridina noctiluca* (hereafter *Cn-luc*) had λ_max_ of 454 nm with a methionine at 178 (M-178-M) while mutants had lower λ_max_ (M-178-R at 435 nm; M-178-K at 447 nm), although these sequences also had other mutations at sites besides 178 (45). Site 178 also varies in c-luciferases of living species (Fig. 1), and notably all *Photeros*, the genus with green-shifted λ_max_. In addition to site 178, variation at sites 280, 375, and 404 shows significant correlation with λ_max_ (Fig. 1; Table 1). Our ANOVA showed a significant interaction between sites 375 and 404, which may be interpreted as epistatic interactions (46).

**Table 1.** Sites significantly associated with phenotypes and their inferred processes of molecular evolution based on estimated rates of synonymous to non-synonymous substitution ratios.

| Phenotype | Site (Cypridina) | Site (Aligned) | ANOVA p | Process | Hyphy p |
| --- | --- | --- | --- | --- | --- |
| Lambda-max<br>(color) | 178 | 207 | 2.2 e-16 | Neutral | NS |
|  | 280 | 311 | 2.5 e-06 | Neutral | NS |
|  | 375 | 406 | 0.013 | Neutral | NS |
|  | 404 | 435 | 0.011 | Purifying | 0.007 |
|  | 375:404 | 406:435 | 0.013 | Epistasis | NA |
|  | 74 | 102 | 0.0001 | Positive | 0.0906 |
|  | 114 | 142 | 0.0033 | Positive | 0.0015 |
|  | 160 | 189 | 0.0053 | Positive | 0.0509 |
| Light decay rate<br>(kinetics) | 19 | 41 | 0.0002 | Positive | 0.0256 |
|  | 160 | 189 | 7.99E-05 | Positive | 0.0509 |
|  | 232 | 261 | 0.003 | Positive | 0.0005 |

### c-Luciferase and the diversification of function

Comparison of c-luciferases and bioluminescence phenotypes shows how multiple evolutionary forces acted on one gene, diversifying courtship signals. We first examined patterns of non-synonymous to synonymous (dn:ds) rates of mutation in the four sites from mutagenesis experiments that most strongly influence λ_max_. Three of these sites (178, 280, 375) failed to reject the null model of neutral evolution. One site (404) had low dn:ds (p=0.071) in the FEL analysis of hyphy. Each of these sites changed λ_max_ in mutagenesis experiments and three of them (280, 375, 404) are fixed within courtship-signaling cypridinids, fixed within non-courtship species, but different between courtship-signaling and non-courtship cypridinids.

In addition to mutated sites, we checked sites under positive selection (dn>ds) for significant associations with biochemical functions. Of the thirteen sites under episodic diversifying selection, five are significantly correlated to bioluminescence phenotypes. Site 160 is under diversifying (positive) selection and correlated significantly with both λ_max_ and light decay of c-luciferases (Fig. 1; Table 1). Four additional sites under positive selection are also correlated with function: sites 74 and 114 are correlated with λ_max_ and sites 19 and 232 are correlated with rates of light decay. These results show how different processes of molecular evolution, including positive and purifying selection, and neutral evolution, together contributed to diversification of biochemical phenotypes that may be important for cypridinid signals (Fig. 4).

**Figure 4.**
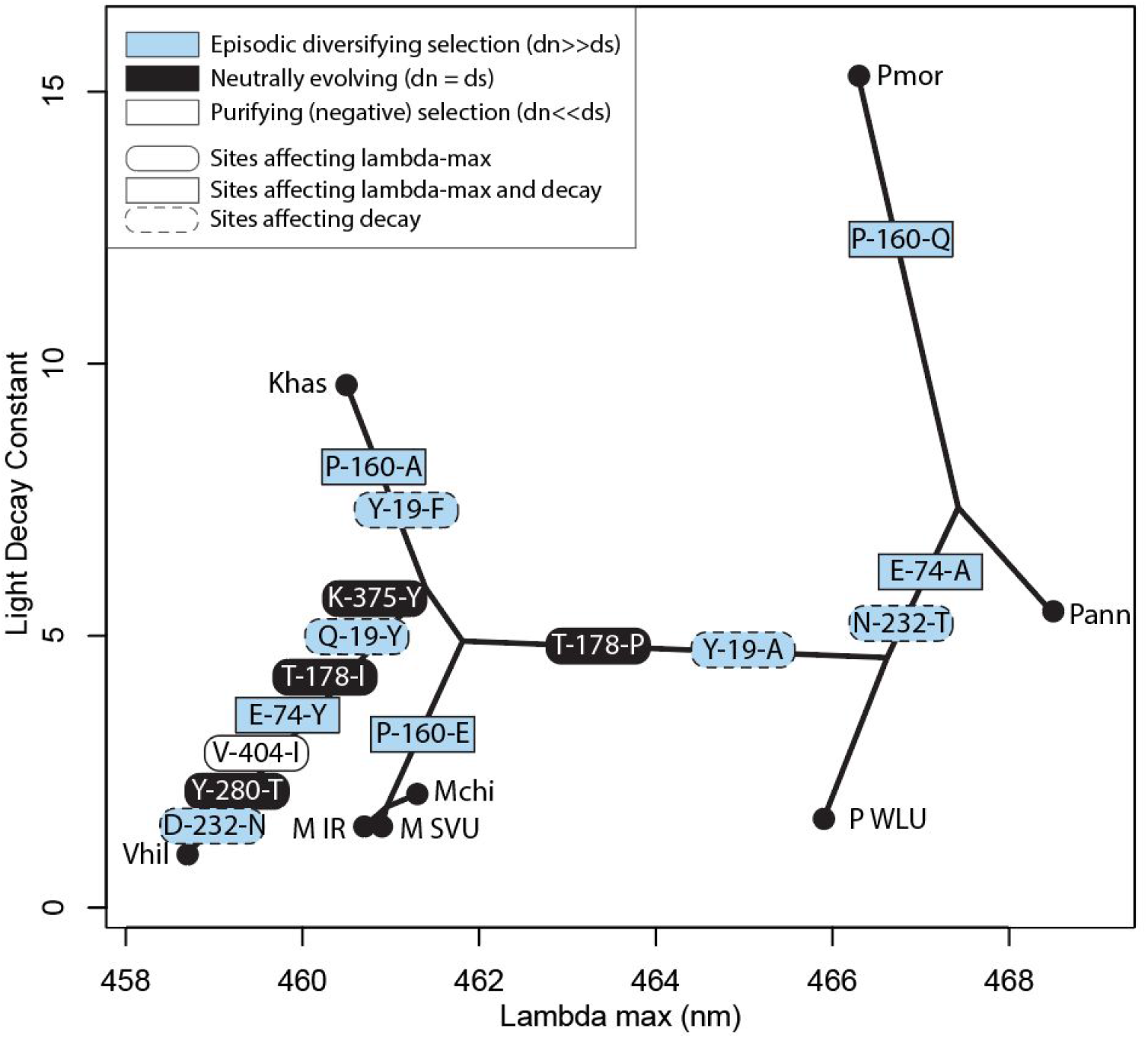
Bioluminescent phenotypes, including *λ_max_* (“Lambda max” on the x-axis) of emission spectra and decay of light emission (“Light decay constant on the y-axis), diversified during the evolution of Cypridinidae, perhaps through different mutations in c-luciferases, and influenced by different evolutionary processes. Circles represent points that different species for which we have (N = 8) occupy in this phylophenospace of bioluminescence, connected by branches representing phylogenetic relationships. Shapes and colors on branches represent inferred mutations and evolutionary processes acting on c-luciferases and correlated with bioluminescence phenotypes. Rectangular sites are correlated with both decay and color, sites with rounded edges either affect color (solid border) or correlate with decay (dashed border). The color of the shapes represent evolutionary processes inferred at those sites, blue is positive selection (dn≫ds), white is purifying selection (dn≪ds), and black is neutral evolution. Each site has a number corresponding to the homologous site in *Cypridina noctiluca* and two single-letter abbreviations for amino acids representing evolved changes at that site. Species abbreviations are Vhil = *Vargula hilgendorfii*, Khas= *Kornickeria hastingsi*, M IR = *Maristella sp*. IR, M SVU = *M. sp*. SVU, Mchi = *M. chicoi*, P WLU = *Photeros sp* WLU, Pann = *P. annecohenae*, Pmor = *P. morini*.

## Discussion

The genetic basis of species diversification remains understudied because many species differences involve many genetic differences across species. At the same time, scientists know very few cases where a single gene contributes to important and extensive phenotypic variation across radiations of species - which could provide tractable systems to study the molecular evolution of diversification. The bioluminescent signals of cypridinid ostracods provide a prime example of signals that vary in multiple parameters across species, and here we show signal diversity to be impacted in important ways by at least a single gene, c-luciferase. Only two previously published c-luciferase genes existed in the literature and we report multiple new luciferase genes and biochemical phenotypes from diverse cypridinid species. Our results indicate various evolutionary forces acted on c-luciferase, contributing to diversification of phenotypes. In addition to some sites in c-luciferase that control color evolving neutrally or under purifying selection, we found episodic diversifying selection on luciferases at amino acid sites strongly associated with changes in both color and enzyme kinetics. In addition to connecting specific mutations to the diversification of biochemical functions, it is interesting to hypothesize how organismal and biochemical phenotypes relate to each other during evolution of this system. As with most biological phenomena, variation in phenotypes could be directly under selection, non-functional, and/or influenced by phylogenetic or biochemical constraints.

First, differences in molecular phenotypes could be shaped by natural and/or sexual selection. Patterns of variation in c-luciferase sequences support selection as one driver of phenotypic variation of biochemical properties of light production. We found multiple sites (19, 160, and 232) in c-luciferases evolving under episodic diversifying selection are also correlated significantly with the rate of decay of light, suggesting selection diversified enzyme kinetics. Of particular interest is site 160 because it is one of only a handful of sites that differs between *Photeros annecohenae* and *P. morini*. Despite strong similarity in c-luciferases in these two species, their enzymes have dramatic differences in measures of decay rate (Fig. 1) that are related to duration of courtship pulses (20). However, which selective forces influence rates of light decay at the organismal level is not yet certain because there are very few experiments on the fitness effects of different ostracod signals (47). We hypothesize pulse duration, in part dictated by enzyme kinetics (20), may be important for fitness via inter-specific anti-predator displays and/or through mate recognition or choice. Consistent with this hypothesis, there is extensive variation among species, with pulse durations in courtship signals ranging from ~0.15 - 9 seconds across species (20, 24). In addition to sexual selection, c-luciferase kinetics could also be under selection at the organismal level due to changes in environmental factors like temperature, pH, or salinity. Because cypridinid luciferase is secreted externally, and bioluminescent ostracods are globally distributed, environmental factors affecting enzyme function vary widely. Indeed, previous *in vitro* expression of *C. noctiluca* luciferase showed temperature-dependent differences in activity (32, 48). Increased sampling of taxa living in different temperatures, including deep-sea cypridinids, could allow testing of varying conditions and the role that adaptation may play in constraining or facilitating changes in rates of light decay.

Another possibility is that variation in some bioluminescent functions are selectively neutral, which seems to be true for color, including emission width (FWHM) and perhaps *λ_max_*. We see no clear pattern in variation of FWHM, and no correlation of this parameter with any positively selected sites in c-luciferases. Although we do not yet have good candidate mutations for the genetic basis of FWHM, we do have such candidates for *λ_max_*, thanks to mutagenesis experiments (45). One site (178) strongly affects *λ_max_* and has a dn:ds ratio indistinguishable from one, consistent with a minimal impact on fitness, suggesting it evolved neutrally. In contrast, three sites (74, 114, 160) are correlated with changes in λ_max_ and under positive selection, which could indicate selection drove some changes in color. The patterns of amino acid differences in these selected sites also show differences between non-courtship and courtship-signaling species. At the organismal level, λ_max_ is non-random with respect to phylogeny (Fig. 1, 3) and we observe evolutionary shifts in λ_max_ between non-courtship and courtship-signaling species and separately, along the branch leading to *Photeros*. If the color change between non-courtship and courtship-signaling species is robust to greater taxon sampling, it could be adaptive, perhaps linked to environmental differences or differences in predators’ vision. In contrast, the color change in *Photeros* does not seem adaptive. Because *Photeros* are among the few signaling species that live in seagrass (22), the “reflectance hypothesis” (49) could predict the ancestor of *Photeros* underwent an adaptive shift to a greener signal to increase signal efficacy, with a correlated shift to grass bed habitat. However, even *Photeros* species that live in non-grass habitats (like *P. morini*) have green-shifted λ_max_ (Fig. 1, 3). Furthermore, an adaptive green-shift is not supported by patterns of substitution in c-luciferases because none of the positively selected sites are fixed in *Photeros*, while also different from non-*Photeros*. Finally, we did not find evidence of positive selection specifically along the branch leading to *Photeros*. If *λ_max_* and FWHM are in fact neutral for organismal fitness, other factors like constraints could instead dictate changes we observed in color, especially *λmax*.

Biochemical constraints like pleiotropy are likely to influence the evolution of bioluminescent phenotypes. One site in c-luciferase (160) is under positive selection and correlated to both λ_max_ and light decay. This suggests a possible role of pleiotropy in phenotypic evolution if mutations in c-luciferase affect multiple phenotypic parameters at once. However, counter to a pervasive role for pleiotropy, we do not find a strong relationship between λ_max_ and light decay constant across all species in our study (Fig. S3). Instead, all *Photeros* have similar λ_max_ while rates of light decay vary considerably between those species. Still, we cannot fully rule out biochemical constraint as a driver for the evolution of emission spectra because such constraints may have changed during evolution, for example during the shift in λ_max_ in early *Photeros*. If so, modern *Photeros* could have biochemical constraints that differ from ancestral species, now allowing rates of light decay to change independently of λ_max_. Testing an evolutionary change in biochemical constraint would entail extensive mutagenesis and expression experiments guided by reconstructing the history of *Photeros* luciferase. Constraints may also arise from epistatic interactions among sites, of which we find some evidence in structuring phenotypic differences between species despite lacking statistical power to exhaustively cover all site-by-site interactions. As phenotypes like color and/or decay evolve, previous changes at certain sites (such as 404) will influence the magnitude of new mutations on function. It is also possible that site-specific epistatic interactions changed during the evolution of c-luciferases, like in other proteins (50, 51).

Taken together, our results support the hypothesis that molecular evolution of c-luciferase impacted the diversity of signal phenotypes in the sea firefly radiation of species. The results further illustrate the potential for varied interactions between molecular evolution, pleiotropy, epistasis, and phenotypes during diversification - even when considering a single gene like c-luciferase. When extrapolated to other genes, such as the presumably many genes affecting behavioral phenotypes and in different environments, the combinatorics become astronomical, providing new perspectives on how life became so incredibly diverse. Such a pluralistic view of evolution, incorporating many different processes at different levels of organization (52, 53), allows for a more holistic understanding of how biodiversity originates.

## Materials and Methods

All data and code for all analyses are available on GitHub: https://github.com/ostratodd/Cypridinidae_Emission.

### Specimen Collection

We collected animals using small nets while SCUBA diving, or by setting baited traps, as previously described (54). We collected multiple species from each of four Caribbean locations: Discovery Bay, Jamaica; Bocas del Toro, Panama; South Water Caye, Belize; and Roatan, Honduras. We also analyzed one species from Catalina Island, California and two from Isla Magueyes, Puerto Rico. We purchased dried *Vargula hilgendorfii*, which originates in Japan (Carolina Biological). We report specific collection localities and size measurements (SI, Table S5). For transcriptomes, we preserved specimens in RNALater (Invitrogen). We carried some animals alive to UCSB to measure emission spectra and others we dried using different methods depending on what tools were available on-site. To induce bioluminescence, we crushed live or dried specimens in seawater using a small plastic pestle, usually one animal at a time.

### Luciferase discovery and amplification

We discovered 12 putative luciferase genes from transcriptomes of cypridinid ostracods. To find these, we queried a previously published transcriptome of *V. tsujii* (55), and new transcriptomes using whole bodies of luminous cypridinids. We will use these transcriptomes for a future phylogenetic study, so we present here paired-end transcriptomes containing putative luciferases, submitted to BioProject PRJNA589015. Metadata from sequencing and NCBI accession number are included in (Table S5).

For six species, we amplified putative c-luciferase sequences via PCR and confirmed nucleotide sequences with Sanger sequencing. In many cases, we found complete assembly of putative luciferase genes from transcriptomes depends on which assembly programs and parameters we used. We believe the VWD domains contained in luciferases (38) have duplicated wildly in ostracods, leading to challenges when assembling transcriptomes from multiple individuals. In some cases, the final assembly in our BioProject differs from early assemblies we used to design primers in calling different regions of a single luciferase gene different isoforms. We analyzed the Sanger-sequenced genes and provide a multiple sequence alignment of corresponding sequences (Fig. S4).

### Luciferase Expression In Vitro

We expressed putative c-luciferases from exemplars of four major clades of cypridinids, *Maristella, Photeros, Kornickeria*, and *Vargula*. Using mammalian cells, we expressed three putative c-luciferase proteins (*Maristella sp. SVU, Vargula tsujii* and *Photeros morini*) and in *Pichia* yeast, we expressed three putative luciferases (*V. tsujii* and *M*. sp. “SVU”, and *Kornickeria hastingsi*). We also expressed previously published *Cypridina noctiluca* luciferase (32) in yeast as a positive control. These proteins are secreted into culture media, to which we added luciferin substrate to test the hypothesis that they are luciferases and catalyze a light reaction. For more details, se SI.

### Light catalysis assay

To assess the ability of constructs to catalyze a light reaction, we harvested cell culture media from mammalian or yeast cells from each transfection (approximately 10 - 25 mL per transfection), and concentrated it using 30,000 MWCO centrifugal filters (Amicon), spun from 30-240 minutes at 4,000 x g. After centrifugation, the protein solution was immediately collected for the assay. Varying volumes of concentrated protein solution and luciferin assay mix (Targeting Systems; prepared to manufacturer’s specifications but with unknown concentration for mammalian cells; and for yeast, we procured vargulin from NanoLight Technology (Pinetop, AZ) and suspended it in 10 mM TRIS to a working concentration of 0.01 ng/ul) were added together in a plate reader (Wallac). We measured luminescence in counts per second (CPS) for 10 seconds.

### Phylogeny of luciferases

We generated a gene tree of translated luciferase candidates and closely related genes. We used a published c-luciferase from *V. hilgendorfii* as a query in a similarity search using BLAST, retaining the top 20 hits from each transcriptome searched. In some cases, we did not obtain full-length luciferase transcripts in the assembly created by Trinity (56), which we attribute to polymorphism from pooling individuals. In these cases, we did a second assembly of luciferase fragments using cap3 (57) and selected the longest orfs as putative c-luciferases. We aligned these sequences using MAFFT (58). We used IQTREE (59) to select the best-fit model of protein evolution and to estimate the maximum likelihood phylogeny. We rooted this phylogeny using midpoint rooting to identify putative c-luciferases from new transcriptomes as orthologues to previously published c-luciferases. Both *P. annecohenae* and *P. sp. WLU* had two additional genes (in-paralogs) in this clade, whereas all other species had one direct ortholog of published c-luciferases in the clade. We excluded these two in-paralogs from further analysis because we did not confirm these transcriptome sequences with PCR and because the in-paralog sequence from *P. annecohenae* lacks a signal peptide and is therefore unlikely to be a functional c-luciferase.

### Testing for positive selection in luciferases

We aligned luciferase DNA by codon, first aligning amino acids in MAFFT (58), then matching DNA codons using pal2nal (60). We used Hyphy (61) to compare ratios of synonymous to non-synonymous substitutions. We used MEME and FEL to search for individual codons under diversifying selection (39).

### Emission Spectra

We used a home-built fluorimeter (spectroradiometer) at UCSB to measure emission spectra. We imaged the output orifice of the integrating sphere or a cuvette cell’s transparent wall by a relay lens onto the entrance slit of a spectroradiometer (Acton SpectraPro 300i) equipped with a charge-coupled device camera (CCD) detector (Andor iDus). Because of the limited time-span of bioluminescence, we collected a series of data frames upon introduction of specimens. We usually sampled at 10 time points every two seconds for each emission with 2 seconds integration time, although sample numbers ranged from 5-20, depending on the species. The spectral data acquired by the CCD camera were corrected for instrumental response artifacts by measuring the spectrum of a black body-like light source (Ocean Optics LS-1) and calculating appropriate correction factors. We also took background spectra for each experiment to subtract from the experimental emission spectra (see SI for further details). For the high quality spectra, we employed Savitzky-Golay smoothing using the signal package (62) in R (63) and then calculated λ_max_(the wavelength with the highest emission value) and Full Width Half Max (FWHM, the width of the spectrum in nanometers where emission is half the value of maximum).

### Genetic Basis of Changes in c-luciferase Function

We looked for amino acid changes associated with changes in three functions: λ_max_, FWHM, and light decay constant. To find mutations in luciferase that shift λ_max_, we analyzed previously published data from mutagenesis experiments on *Cypridina noctiluca* luciferase (*Cn-luc*) and data on luciferases from 9 other cypridinid species with both luciferase and emission spectrum data. Kawasaki et al. (45) created 35 variants of *Cn-luc* and measured λ_max_ for each variant plus wild type *Cn-luc*. To measure λ_max_ they added luciferin to *Cn-luc* variants expressed heterologously, finding λ_max_to vary between 435 nm and 463 nm. They did not report complete spectra or FWHM for most of the mutants, instead only reporting λ_max_. Therefore, our subsequent analyses of FWHM use only data from natural c-luciferase sequences and our newly reported FWHM data from emission spectra. For the mutational analysis, each variant contained 1-8 mutated sites compared to wild type, and across all 35 variants, a total of 23 different amino acid sites contained mutations. For the non-*Cn-luc* luciferases, we aligned sequences to *Cn-Luc* using MUSCLE (64) to identify amino acids at sites homologous to those mutated in *Cn-Luc* variants. We refer to sites as numbered for the homologous positions in *Cn-luc*, which differs from their aligned position. For light decay constant, we used previously published data (20).

We analyzed candidate mutations with ANOVA to test for significant associations between variant amino acid sites and functions of c-luciferases, similar to methods of Yokoyama et al. for opsins and absorbances (46). To determine which sites were most highly correlated with each function, we used a model selection approach in the R package MuMIn (ref). We used the dredge function to test combinations of amino acid sites, which formed different models. Dredge sorts models by AIC score. We tallied amino acid sites present in the highest number of best-fit models, and then performed ANOVA using a model with those sites.

## Acknowledgements

We thank C. Pham, T. Halvorsen, Y. Chiu, J. Russo, M. Pfau, and T. Jessup for assistance with cloning and expression. We acknowledge J. Lachat and Y. Nishimiya for assistance with the *Pichia* system and for providing the *C. noctiluca* construct used as a positive control. We are grateful to Z. Terner from the UCSB Statistical Consulting Laboratory for his advice on modeling. We thank K. Niwa for assistance in the early stages of emission spectra collection. We also extend our thanks to Y J. Li, S. Schulz, N. Reda, J. Morin, M D. Ramirez, C. Motta, and V. Gonzalez for assistance with animal collections. Organisms were collected under the auspices of proper permitting from the Jamaican National Environment and Planning Agency (Permit Ref. #18/27), the Belize Fisheries Department (Permit #000003-16), the Honduran Department of Fish and Wildlife (Permit #DE-MO-082-2016), the Puerto Rican Department of Natural and Environmental Resources (DRNA; Permit #2016-IC-113), and Panamanian Ministry of the Environment (MiAMBIENTE; Permit #SE/A-33-17). NMH was funded in part by an NSF EAPSI Fellowship (#1713975), the NSF Graduate Research Fellowship Program, and the UCSB Research Mentorship Program run by Dr. Lina Kim. NSF DDIG #1702011 to EAE provided funding for transcriptomes and emission spectra estimation. JC received funding from the UCSB URCA program. ET was funded by a CSUPERB Research Development Grant for transcriptomes, gene synthesis, and vector cloning. Authors THO, GAG, and ET were funded by NSF (#DEB-1457754, DEB-1457439, and DEB-1457462 respectively.)

## Link to Tables (In Spreadsheet Format)

<u>Link</u> to supplemental figures and legends

Author information (contributions as <u>here</u>):

1. Nicholai Marcus Hensley

- Affiliation: Department of Ecology, Evolution, & Marine Biology, University of California, Santa Barbara
- Contribution: Conceptualization, Funding Acquisition, Methodology, Investigation, Writing - Review & Editing, Project Administration, Data Curation, Formal Analysis, Software
- Competing interests: No competing interests declared
- ORCID: 0000-0003-0681-0808
2. Emily A. Ellis

- Affiliation: Florida Museum of Natural History, University of Florida
- Contribution: Conceptualization, Funding Acquisition, Methodology, Investigation, Writing - Review & Editing
- Competing interests: No competing interests declared
3. Nicole Y. Leung

- Affiliation: Neuroscience Research Institute and Department of Molecular, Cellular and Developmental Biology, University of California, Santa Barbara California
- Contribution: Investigation, Writing - Review & Editing
- Competing interests: No competing interests declared
- ORCID: 0000-0001-8823-1783
4. John Coupart

- Affiliation: Department of Ecology, Evolution, & Marine Biology, University of California, Santa Barbara
- Contribution: Investigation
- Competing interests: No competing interests declared
5. Alexander Mikhailovsky

- Affiliation: Department of Chemistry and Biochemistry, University of California, Santa Barbara
- Contribution: Methodology, Resource, Writing - Review & Editing
- Competing interests: No competing interests declared
6. Daryl A. Taketa

- Affiliation: Department of Molecular, Cellular and Developmental Biology, University of California, Santa Barbara
- Contribution: Methodology, Investigation
- Competing interests: No competing interests declared
7. Michael Tessler

- Affiliation: American Museum of Natural History and New York University, New York
- Contribution: Resources, Writing - Review & Editing
- Competing interests: No competing interests declared
- ORCID: 0000-0001-7870-433X
8. David F. Gruber

- Affiliation: Department of Biology and Environmental Science, City University of New York Baruch College, New York
- Contribution: Funding Acquisition, Resources, Writing - Review & Editing
- Competing interests: No competing interests declared
- ORCID: 0000-0001-9041-2911
9. Anthony W. De Tomaso

- Affiliation: Department of Molecular, Cellular and Developmental Biology, University of California, Santa Barbara
- Contribution: Supervision, Resources
- Competing interests: No competing interests declared
10. Yasuo Mitani

- Affiliation: Bioproduction Research Institute, National Institute of Advanced Industrial Science and Technology (AIST), Sapporo
- Contribution: Funding Acquisition, Supervision, Writing - Review & Editing, Resources
- Competing interests: No competing interests declared
11. Trevor J. Rivers

- Affiliation: Department of Ecology and Evolutionary Biology, University of Kansas
- Contribution: Resources
- Competing interests: No competing interests declared
12. Gretchen A. Gerrish

- Affiliation: Trout Lake Station, Center for Limnology, University of Wisconsin - Madison
- Contribution: Funding Acquisition, Project Administration, Resources, Writing - Review & Editing
- Competing interests: No competing interests declared
- ORCID: 0000-0001-6192-5734
13. Elizabeth Torres

- Affiliation: Department of Biological Sciences, California State University Los Angeles
- Contribution: Funding Acquisition, Resources, Project Administration, Writing - Review & Editing
- Competing interests: No competing interests declared
14. Todd Hampton Oakley

- Affiliation: Department of Ecology, Evolution, & Marine Biology, University of California, Santa Barbara
- Contribution: Conceptualization, Visualization, Writing – Original Draft Preparation, Writing - Review & Editing, Software, Data Curation, Funding Acquisition, Formal Analysis, Supervision, Project Administration, Resources
- Competing interests: No competing interests declared
- ORCID: 0000-0002-4478-915X

