## Supplemental Material for "Molecular evolution of luciferase diversified bioluminescent signals in sea fireflies"

### Supplemental Materials and Methods

*Specimen collection*

We identified most species by the patterns of male courtship signals, targeting each particular display type with a single net and later measuring length and height of individual animals using a dissecting microscope and ocular micrometer. Several of the species, especially from Panama, are undescribed and we refer to those here by field codes, consisting of two or three capital letters. We classify species into genera based on length:height ratio, which is a reliable genus-level characteristic (1, 2). For emission spectra specimen preservation, methods varied by locale. In Roatan and Puerto Rico, we dried ostracods in direct sunlight. In Panama and Belize, we used a drying oven and transported animals in Eppendorf tubes with silica beads as a dessicant.

*Luciferase discovery and amplification*.

We designed luciferase-specific primers (Table S6) to amplify from cDNA and obtain sequences that do not include signal peptides (18 or 19 amino acids from the n-terminus, as inferred with SignalP) because we later used yeast-specific signal peptides during protein expression. we used an initial denaturation of 95°C for 2 min. For 30x cycles, we performed a 95°C denaturation step for 1 min., followed by an annealing phase at varying temperatures per species (*K. hastingsi*: 45.5°C, *P. morini*: 48°C, *M. sp.* SVU: 41.1°C , *M. sp.* SVD: 43.7°C, *M chicoi*: 41.4°C, *V. tsujii*: 45.5°C) for 1:45 min., and then by an extension step at 73°C for 1 min. For *V. tsujii* primers designed from the published transcriptome, we used thermal profile: 40 cycles of 94°C for 35s, 55°C for 30s, 72°C for 1min) and amplified the native signal peptide.

*Luciferase Expression In Vitro*.

We expressed three luciferases in mammalian HEK293 cells. To construct a *V. tsujii* luciferase (VtL) expression vector, we first amplified VtL using primers with engineered restriction sites to clone into a pCR4-TOPO vector. We next excised VtL-pCR4 with XhoI and EcoRI (Promega), and subcloned into a modified pCMV3B mammalian expression vector with a C-terminal mCherry reporter (mCherry-C). The luciferase genes of *P. morini* and *M. sp. SVU* were synthesized and cloned into the mCherry-C vector by Genscript (Piscataway, New Jersey, USA) with flanking restriction enzyme sites. We planned to use mCherry to quantify the concentration of expressed luciferase, but we found high autofluorescence of cell media and/or other secreted proteins to preclude this use. We first transformed cloned constructs into competent *E. coli* cells using the One Shot Chemical Transformation Kit (Invitrogen), and cultured for 24 hours in standard lysogeny broth (LB) with 0.1% kanamycin at 37°C. We verified construct transformation using the engineered restriction enzyme sites in digests and compared them to their expected product size. We extracted these plasmids using the FastPlasmid Mini kit (Qaigen) and assessed concentrations with the Qubit high standard DNA kit (Qubit). For transfection, we cultured mammalian HEK293 cells in Dulbecco's modified Eagle's medium (DMEM), supplemented with 10% fetal bovine serum (FBS) and penicillin/streptomycin (P/S) at 37°C with 5% $CO_2$. We then plated 5 x $10^4$ cells in each well of a 24-well plate one to three days before transfection. Cell medium was changed to DMEM without serum and antibiotics before transfection. We transfected cells with 0.5 µg of vector using Lipofectamine 2000 (Invitrogen), performed according to the manufacturer's instructions. After 4 hours of incubation, we replaced the transfection medium with DMEM+FBS+P/S and allowed the cells to recover for 24 hours. We collected cells via trypsin digestion and reseeded them into 10mL of DMEM+FBS+P/S+1% G418 to select against untransfected cells in 90cm cell plates. We cultured the transfected cells for 3 to 5 days before harvesting and using in light catalysis assays.

For expression in *Pichia* yeast, we cloned sequences into the pPICZ-αC vector at the XhoI and NotI sites following standard procedures (Invitrogen Easy Select *Pichia* kit). First, we analyzed predicted c-luciferases for the presence of a signal peptide at the n-terminal end using SignalP v4.1 (3). We then designed primers for cloning and protein expression to amplify the entire c-luciferase sequence without the native signal peptide, beginning usually 51-54 bp inside the 5' end from the predicted start codon. 3' end primers excluded the native stop codon so that a fusion construct could be generated. Fusion constructs were made via the EasySelection Pichia expression kit (Invitrogen) using the pPICZ-αC vector according to the manufacturer's instructions. Briefly, we used the 5' XhoI site in order to generate fusion c-luciferases with an alpha secretion signal from yeast. We reconstructed the Kecx2 cleavage site with one Glycine Alanine repeat region via PCR. On the 3' end, we used NotI; this would result in the addition of extra amino acids in our expressed proteins on the c-terminal end before inclusion of the fusion c-myc epitope and histidine tags. We transformed newly-made, linearized constructs into *Pichia* using electroporation with a BioRad Micro-Pulser using the Sc2 program (1.5 V). After electroporation, *Pichia* were allowed to recover in selective media for an hour before plating. We initially selected for recombinant *Pichia* colonies using two concentrations of zeocin (100 and 500 mg/mL). After three days of growth, individual colonies were replica-plated at high zeocin concentrations (1,000 and 2,000 mg/mL) to try and screen for high copy-number integrants for our gene of interest. After one day, we selected single colonies that grew best at high zeocin concentrations to induce protein expression according to the manufacturer's guidelines. To stabilize the pH of the media for extended expression, colonies were grown in 25mL buffered media with glycerol in baffled flasks until the $OD_{600}$ reached 2.0 - 8.0. For our colonies, this occurred after 72 hrs due to suboptimal shaking conditions. We then calculated the amount of original growth we would need for an $OD_{600}$ of 1.0 in 30mL expression media, spun down the appropriate volume of the original colonies at 3,000 g for 5min., removed the glycerol media, and resuspended the pellet in 30mL of buffered media with methanol in a 125 mL baffled flask. Flasks were shaken in a table-top incubator at 29.5C at 300 rpm for 3 days, with media supplemented with 100% methanol every 24hrs to maintain a 0.5% volume of methanol in culture.

*Emission Spectra*

In earlier trials of emission spectra data collection, we introduced specimens into a test tube placed inside Spectralon-coated 150 mm diameter integrating sphere (Labsphere). But because we report relative rather than absolute levels of light at different wavelengths, we abandoned the integrating sphere in later trials for a rectangular quartz cuvette to increase sensitivity of our analyses.

To correct for variation in emission, background noise, and quality during data collection, we first summed all collection time points for one sample, then subtracted each background value for each wavelength, and corrected using measurements from the black body radiator before standardizing each spectrum, setting the maximum value to 1.0 as the wavelength with the most photons. Some specimens did not yield strong light emission, probably due to variation in drying. We filtered low quality data using signal to noise ratio. Specifically, we sorted emission values at each wavelength from lowest to highest and averaged the lowest 1000 data points to estimate a baseline emission value ($E_{min}$). We then found the maximum emission value ($E_{max}$). From this, we calculated a signal to noise value as $E_{min}/E_{max}$, removing trials where $E_{min}/E_{max} < 0.02$.

#### Supplemental Figures and Tables

**Supplemental Figure S1** - Maximum likelihood phylogeny of sequences most similar to published c-luciferases.

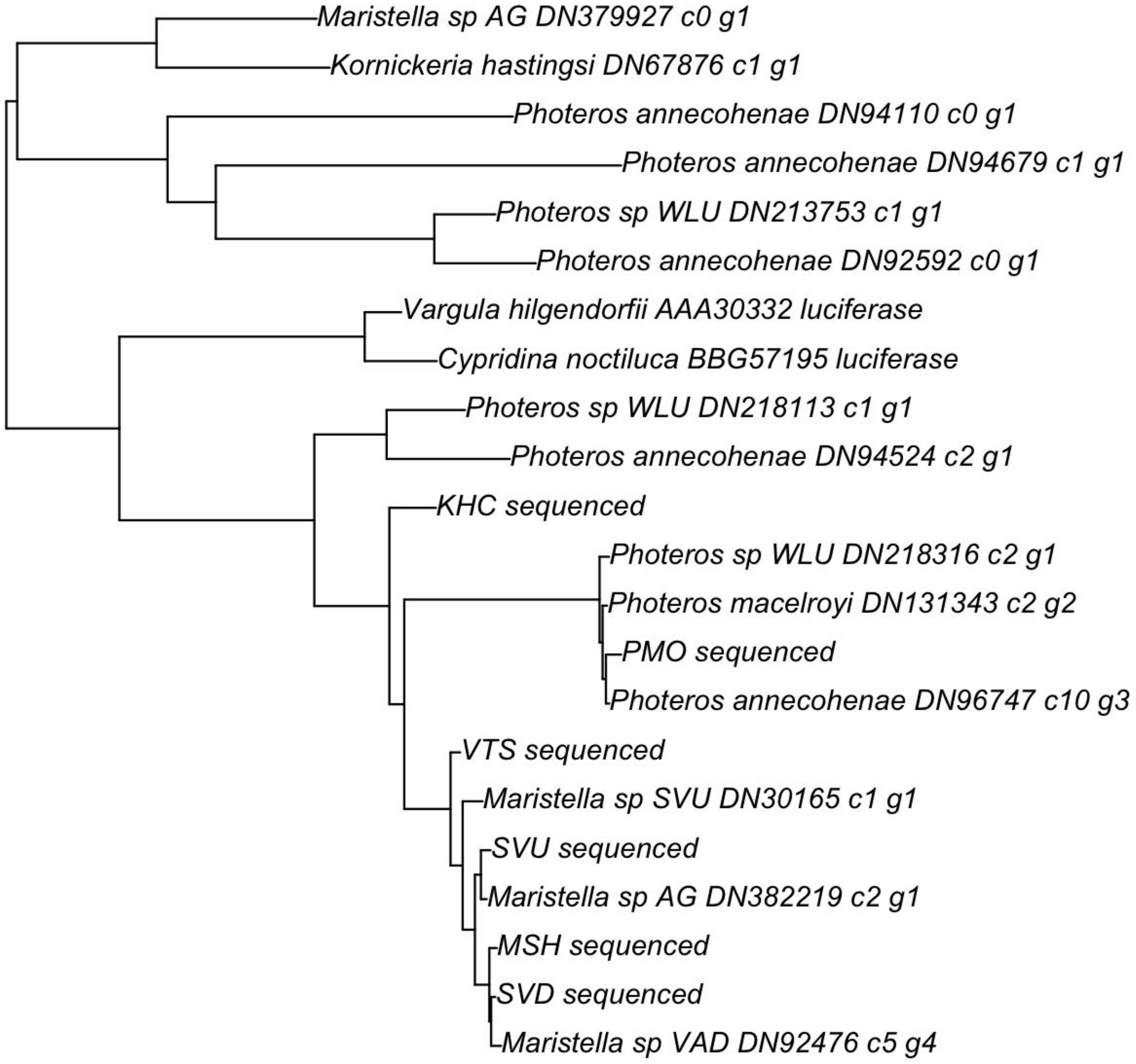

**Supplemental Figure S2**. Full Width of emission spectrum at Half of the Maximum value (FWHM) in nanometers (nm) from new emission spectra from 20 species and previously published spectra from 3 species.

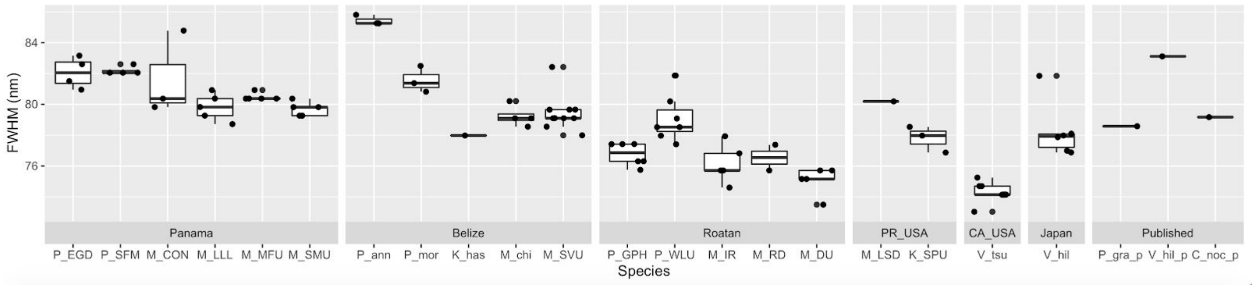

**Supplemental Figure S3**. We find no relationship between the color of light emission and light decay constants (Pearson's correlation test, t = , df = 16, p = 0.4). Scatter plot of the average decay (x-axis) and peak emission spectra (y-axis) measured for different species of luminescent ostracod (N = 17 species). Data are colored by genus, and shape by country of origin. Although *Photeros* greatly differ in their peak emission spectra than other genera, there is no strong pattern for differences in light decay constant.

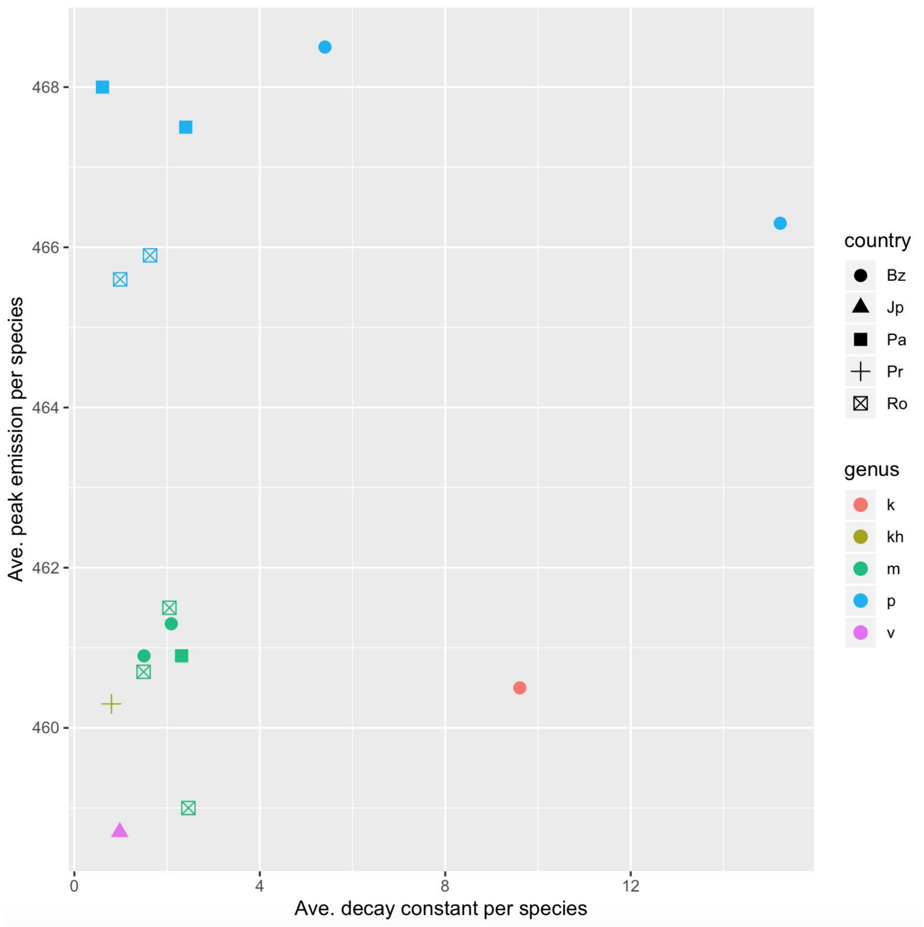

**Supplemental Figure S4**. Multiple sequence alignment for putative and functional c-luciferase sequences

```
Vargula_hilgendorfii      ----------------------MKIIILSVILAYCVTDNCQDACPVEAEP    28
Cypridina_noctiluca_      ----------------------MKTLILAVALVYCVTVNCQE-CPYVADP    27
Kornickeria_hastings      -------------------------------------KDCFESSFHSLE-    12
Vargula_tsujii_seque      -------------------------------------QDCYESTWASND-    12
Maristella_chicoi_se      -------------------------------------EDCYEFTWASND-    12
Maristella_sp_SVD_se      ----------------------------------------YEFTWASND-    9
Maristella_sp_SVU_se      -------------------------------------QDCYEFTWASND-    12
Maristella_sp_AG_DN3      MGLKTHSAFVISREIRSGIMWLQSLL-LLAGICFCAGQDCYEFTWASND-    48
Maristella_sp_VAD_DN      -------------------MWLQSLL-LLAGICFCAGEDCYEFTWASND-    29
Photeros_sp_WLU_DN21      -------------------MWLQNLLILLVGIYFSSGEECAQTSPTYLD-    30
Photeros_macelroyi_D      -------------------MWLQNLLILLVGIYFSSGEECAQTSLTSLD-    30
Photeros_morini_sequ      -------------------------------------QECAQTSLTSLD-    12
Photeros_annecohenae      -------------------MWLQNLLILLVGIYFSSGEECAQTSLTSLD-    30

Vargula_hilgendorfii      PSSTPTVPTSCEAKEGECIDTRCATCKRDILSDGLCENKPGKT--CCRMC    76
Cypridina_noctiluca_      PN---TVPTSCEAKEGECIDSSCSTCTRDILSDGLCENKPGKT--CCRMC    72
Kornickeria_hastings      ------FPSSCEAQNGICIDSECKDCNEVMFGDGLCENAGGASPKCCRDC    56
Vargula_tsujii_seque      ------YPSSCEALNGRCVDSACGSCDEVLFGDGLCENAGGASPKCCRDC    56
Maristella_chicoi_se      ------YPSSCEAINGRCVDSACGSCDQVLFGEGLCENAGGASPKCCRDC    56
Maristella_sp_SVD_se      ------YPSSCEAINGRCVDSACGSCDQVLFGDGLCENAGGASPKCCRDC    53
Maristella_sp_SVU_se      ------FPSSCEAINGRCVDSACGSCDEVLFGDGLCENAGGASPKCCRDC    56
Maristella_sp_AG_DN3      ------FPSSCEAINGRCVDSACGSCDEVLFGDGLCENAGGASPKCCRDC    92
Maristella_sp_VAD_DN      ------YPSSCEAINGRCVDSACGSCDQVLFGDGLCENAGGASPKCCRDC    73
Photeros_sp_WLU_DN21      -----SVPSSCEAQRGVCKDSSCSGCDKALYGDDMCENSGVANAKCCRDC    75
Photeros_macelroyi_D      -----SVPSSCEAQRGVCKDSSCSGCDKALYGDDMCENSGIDSAKCCRDC    75
Photeros_morini_sequ      -----SAPSSCEAQKGVCKDPSCSGCDKALYGDDMCENSGVANAKCCRDC    57
Photeros_annecohenae      -----SVPSSCEAQKGVCKDSSCSGCDKALYGDDMCENSGVANAKCCRDC    75

Vargula_hilgendorfii      QYVIECRVEAAGYFRTFYGKRFNFQEPGKYVLARGTKGGDWSVTLTMENL    126
Cypridina_noctiluca_      QYVIECRVEAAGWFRTFYGKRFQFQEPGTYVLGQGTKGGDWKVSITLENL    122
Kornickeria_hastings      PEIVRCRASAAGFFTTFYGKKFNLQEPGTYLLSEDCVGGLWSLYVTLVNI    106
Vargula_tsujii_seque      PEIVRCRVSAAGYFQTFYGQRFNLQVPGTYLLSEDCVGGLWSLYVNLANI    106
Maristella_chicoi_se      PEIVRCRASAAGFFHTFYGQRFNLQVPGTYLLSEDCVGGLWSLYVNLANV    106
Maristella_sp_SVD_se      PEIVRCRASAAGFFHTFYGQRFNLQVPGTYLLSEDCVGGLWSLYVNLANV    103
Maristella_sp_SVU_se      PEIVRCRASAAGFFHTFYGQRFNLQVPGTYLLSEDCVGGLWSLYVNLANI    106
Maristella_sp_AG_DN3      PEIVRCRASAAGFFHTFYGQRFNLQVPGTYLLSEDCVGGLWSLYVNLANI    142
Maristella_sp_VAD_DN      PEIVRCRASAAGFFHTFYGQRFNLQVPGTYLLSEDCVGGLWSLYVNLANV    123
Photeros_sp_WLU_DN21      PEVVRCRAAGAGYFTTFYGKRFNFQVPGKYLLSEDCVGGLWSLYVNLAPI    125
Photeros_macelroyi_D      PEVVRCRAAGGGYFTTFYGKRFNFQVPGKYLLSEDCVGGLWSLYVNLAPI    125
Photeros_morini_sequ      PAVVKCRAAGAGYFTTFYGKRFNFQVPGKYLLSEDCVGGLWSLYVNLAPI    107
Photeros_annecohenae      PAVVKCRAAAAGYFTTFYGKRFNFQVPGKYLLSEDCVGGLWSLYVNLAPI    125

Vargula_hilgendorfii      DGQKGAVLTKTTLEVAGDV-IDITQATADPITVNGGADPVIANPFTIGEV    175
Cypridina_noctiluca_      DGTKGAVLTKTRLEVAGDI-IDIAQATENPITVNGGADPIIANPYTIGEV    171
Kornickeria_hastings      AGEKGAVLGSVKM-IVGEVTVDIVQKGG-PVTVNGGSVAIDSNPFSIGDV    154
Vargula_tsujii_seque      EGEKGAVLDSVKM-VVGDVTVDIKQKVG-AVTVNGGSVEIDSNPFSIGDV    154
Maristella_chicoi_se      EREKGAVLDSVKM-VVGDVTLDIKQKVG-SITVNGGTVDIDSNPFSIGDV    154
Maristella_sp_SVD_se      EGEKGAVLDSVKM-VVGDVTLDIKQKVA-SITVNGGTVDIDSNPFSIGDV    151
Maristella_sp_SVU_se      EGEKGAVLDSVKM-VVGDVTLDIKQKVG-SITVNGGSVEIDSNPFSIGDV    154
Maristella_sp_AG_DN3      EGEKGAVLDSVKM-VVGDVTLDIKQKVG-SITVNGGSVAIDSNPFSIGDV    190
Maristella_sp_VAD_DN      EGEKGAVLDSVKM-VVGDVTLDIKQRLG-YITVNGGAVAIDSNPFSIGDV    171
Photeros_sp_WLU_DN21      EGQKGAALESVNM-IVGDVTVDITQKGG-PIVVNKGDVPIDSNPFSIGDV    173
Photeros_macelroyi_D      EGQKGAALESVNM-IVGDVTVDITQKGG-PIVVNKGDVPIDSNPFSIGDV    173
Photeros_morini_sequ      EGQKGAALESVNM-IFGDVTIDITQKGG-PIVVNKGDVQIDSNPLSIGDV    155
Photeros_annecohenae      EGQKGAALESVNM-IVGDVTVDITQKGG-PILVNKGDVPIDSNPFSIGDV    173

Vargula_hilgendorfii      TIAVVEIPGFNITVIEFFKLIVIDILGGRSVRIAPDTANKGLISGICGNL    225
Cypridina_noctiluca_      TIAVVELPGYNITVIEFFKLIVIDILGGRSVRIAPDTANKGMISGLCGDL    221
Kornickeria_hastings      TIAIVHTPNFDVAVIEFLKLVTFDIIHGRAFRLAPDFLYRDRTCGLCG-V    203
Vargula_tsujii_seque      TIAIVHTPNFDVSVIEFLKLVTFDILQGRAFRLAPDFLYADRTCGLCG-V    203
Maristella_chicoi_se      TIAIVHTPYFDVSVIEFLKLVTFNILQGRAFRLAPDFLYADRTCGLCG-V    203
Maristella_sp_SVD_se      TIAIVHTPFFDVSVIEFLKLVTFDILQGRAFRLAPDFLYADRTCGLCG-L    200
Maristella_sp_SVU_se      TIAVVYTPYFSVSVIEFLKLVTFDILQGRAFRLAPDFLYADRTCGLCG-L    203
Maristella_sp_AG_DN3      TIAVVYTPYFSVSVIEFLKLVTFDILQGRAFRLAPDFLYADRMCGLCG-L    239
Maristella_sp_VAD_DN      TIAIVHTPYFDVSVIEFLKLVTFDILQGRAFRLAPDFLYADRTCGLCG-L    220
Photeros_sp_WLU_DN21      TIAVVTPPFSHVSVIEFRRIVTFDIISGRAFRLAPDFYYRDRTCGLCG-L    222
Photeros_macelroyi_D      TIAIVTPPFSKVSVIEFLRIVTFDIISGRAFRLAPDFYYRDRTCGLCG-L    222
Photeros_morini_sequ      TIAIVTPPFSKVSVIEFLRIVTFDIISGRAFRLAPDFYYRDRTCGLCG-L    204
Photeros_annecohenae      TIAIVTPPFSKVSVIEFLRIVTFDIISGRAFRLAPDFYYRDRTCGLCG-L    222

Vargula_hilgendorfii      EMNDADDFTTDADQLAIQPNINKEFDGCPFYGNPSDIEYCKGLMEPYRAV    275
Cypridina_noctiluca_      KMMEDTDFSSDPEQLAIQPKINQEFDGCPLYGNPEDITYCKGLLEPYKDS    271
Kornickeria_hastings      MSNEPTDFIDNPDQLAVQDKINKDIDGCPLSGNPSDVEYCKNKMQPYKVA    253
Vargula_tsujii_seque      MSDEPTDFIDNPDQLAIQDQMNQDIDGCPLSGNPSDVEYCVNKMQPYKDG    253
Maristella_chicoi_se      MSDEPSDFIDNPDQLAIQDLMNQDVEGCPLSGNPSDIEYCVNKMQPYKDG    253
Maristella_sp_SVD_se      MSDEPSDFIDNPDQLAIQDLMNQDVEGCPLSGNPSDIEYCVNKMQPYKDG    250
Maristella_sp_SVU_se      MSDEPSDFIDDPDQLAIQDIMNQDVEGCPLSGNPSDAEYCLNKMQPYKDG    253
Maristella_sp_AG_DN3      MSDVPSDFIDNPDQLAIQDIMNQDVEGCPLSGNPSDVEYCLNKMQPYKDG    289
Maristella_sp_VAD_DN      MSDEPSDFIDNPDQLAIQDLMNQDVEGCPLSGNPSDIEYCVNKMQPYKDG    270
Photeros_sp_WLU_DN21      MTTEASDFVNNPDQLAIKDQIDQDVEGCPLSGNPSDVEYCTKYLKPHQDN    272
Photeros_macelroyi_D      MTTEASDFVNCPDQLAIKDQIDQDVEGCPLSGNPSDVEYCTKYLKPHQDN    272
Photeros_morini_sequ      MTTEASDFVNCPDQLSIKDQIDQDVEGCPLSGNPSDVEYCTKYLKPHQDN    254
Photeros_annecohenae      MTTEASDFVNCPDQLAIKDQIDHDVEGCPLSGNPSDVEYCTKYLKPHQDN    272
```

```
Vargula_hilgendorfii   CRN--NINFYYYTLSCAFAYCMGGEERAKHVLFDYVETCAAPETRGTCVL   323
Cypridina_noctiluca_   CRN--NINFYYYTLSCAFARCMGGDERASHVVLDYRETCAAPETRGTCVL   319
Kornickeria_hastings   CINYNDVNFATYLYSCALAYCMGGEDRVEDVIFDYVEACVEPITRATCVM   303
Vargula_tsujii_seque   CVNKNDVHFGTYLYACALAYCMGGDDRVEDVIFEYAEACVEPIGRATCVM   303
Maristella_chicoi_se   CINKNDVHFGTYLYACALAYCMGGDDRVEDVIFEYAEACVEPIGRATCVM   303
Maristella_sp_SVD_se   CINKNDVHFGTYLYACALAYCMGGDDRVEDVIFEYAEACVEPIGRATCVM   300
Maristella_sp_SVU_se   CINKNDVHFGTYLYACALAYCMGGDDRVEDVIFEYAEACVEPIGRATCVM   303
Maristella_sp_AG_DN3   CINKNDVHFGTYLYACALAYCMGGDDRVEDVIFEYAEACVEPIGRATCVM   339
Maristella_sp_VAD_DN   CINKNDVHFGTYLYACALAYCMGGDDRVQDVIFEYAEACVEPIGRATCVM   320
Photeros_sp_WLU_DN21   CNNGDAIHFATYVYACALAYCMGGDDRAEDVAMDYQEACVDPIGRGTCVM   322
Photeros_macelroyi_D   CKNGDAIHFATYVYACALAYCMGGDDRAEDVAMDYQEACVDPIGRGTCVM   322
Photeros_morini_sequ   CKNGDAIHFATYVYACALAYCMGGDDRAEDVAMDYQEACVDPIGRGTCVM   304
Photeros_annecohenae   CKNGDAIHFATYVYACALAYCMGGDDRAEDVAMDYQEACVDPIGRGTCVM   322

Vargula_hilgendorfii   SGHTFYDTFDKARYQFQGPCKEILMAADCYWNTWDVKVSHRDVESYTEVE   373
Cypridina_noctiluca_   SGHTFYDTFDKARYQFQGPCKEILMAADCYWNTWDVKVSHRNVDSYTEVE   369
Kornickeria_hastings   NGHTYYDTFDKTSYQFQAPCK-VLFAKDCAGDEWEVTITHKAAGTYTEVE   352
Vargula_tsujii_seque   NGHTYYDTFDKTSYQFQAPCK-VLFAKDCTDDDWEVTITHKAVGEYTEVE   352
Maristella_chicoi_se   NGHTYYDTFDKTSYQFQAPCK-VLFAKDCTDDDWEVTITHKAAGEYTEVE   352
Maristella_sp_SVD_se   NGHTYYDTFDKTSYQFQAPCK-VLFAKDCTDDDWEVTIVHKAAGEYTEVE   349
Maristella_sp_SVU_se   NGHTYYDTFDKSSYQFQAPCK-VLFAKDCTDDEWEVTITHKEAGEYTEVE   352
Maristella_sp_AG_DN3   NGHTYYDTFDKTSYQFQAPCK-VLFAKDCTDDEWEVTITHKAAGEYTEVE   388
Maristella_sp_VAD_DN   NGHTYYDTFDKTSYQFQAPCK-VLFAKDCTDDEWEVTIVHKAAGEYTEVE   369
Photeros_sp_WLU_DN21   SGHTFYDTFDKTSFQYQAPCK-VQFSKDCIGDDWEVSITYKPKADYTVVD   371
Photeros_macelroyi_D   SGHTFYDTFDKTSFQYQTPCK-VQFSKDCIGDDWEVSITYKPKADYTVVD   371
Photeros_morini_sequ   SGHTFYDTFDKTSFQYQTPCR-VQFSKDCVGDDWEVSITYKPKADYTVVD   353
Photeros_annecohenae   SGHTFYDTFDKTSFQYQTPCK-VQFSKDCIGDDWEVSITYKPKADYTVVD   371

Vargula_hilgendorfii   KVTIRKQSTVVDLIVDGKQVKVGGVDVSIPYSSENTSIYWQDGDILTTAI   423
Cypridina_noctiluca_   KVRIRKQSTVVELIVDGKQILVGGEAVSIPYSSQNTSIYWQDGDILTTAI   419
Kornickeria_hastings   KVTVRYFQTLIDLISEGKQVLVNGTEVSVPYNKGDTSIYMYD-NLITTAV   401
Vargula_tsujii_seque   KVTVRYFQTLIDLISETKKVFVNGTEVSVPYNYGDTSIYMYD-NLITTAV   401
Maristella_chicoi_se   KVTVRYFQTLIDLIAESKKVFVNGTEVSVPYNYGDTSIYMYD-NLITTAV   401
Maristella_sp_SVD_se   KVTVRYFQTLIDLVAENKKVFVNRTEVSVPYNYGDTSIYMYD-NLITTAV   398
Maristella_sp_SVU_se   KVTVRYFQTLIDLVAENKKVFVNGTEVSVPYNYGDTSIYMYD-NLVTTAV   401
Maristella_sp_AG_DN3   KVTVRYFQTLIDLIAENKKVFVNGTEVSAPYNYGDTSIYMYD-NLITTAV   437
Maristella_sp_VAD_DN   KVTVRYFQTLIDLVAESKKVFVNGTEVSVPYNYGDTSIYMYD-NLVTTAV   418
Photeros_sp_WLU_DN21   KVTVRYFATLIDLIPEGRQVLVNGSAVSVPFNYADTSVYMYE-NLITTAV   420
Photeros_macelroyi_D   KVTVRYFATLIDLIPEGRQVLVNGSAVSVPFNYADTSIYMYE-NLITTAV   420
Photeros_morini_sequ   KVTVRYFATLIDLIPEGRQVLVNGSAVSVPFNYADTSIYMYE-NLITTAV   402
Photeros_annecohenae   KVTVRYFATLIDLIPEGRQVLVNGSAVSVPFNYADTSIYMYE-NLITTAV   420

Vargula_hilgendorfii   LPEALVVKFNFKQLLVVHIRDPFD-GKTCGICGNYNQDSTDDFFDAEGA-   471
Cypridina_noctiluca_   LPEALVVKFNFKQLLVVHIRDPFD-GKTCGICGNYNQDFSDDSFDAEGA-   467
Kornickeria_hastings   LPGAVVVKYNFEQMLALHIRDPYE-RRSCGLCGIWDLDKSNDGPDNQYVD   450
Vargula_tsujii_seque   LPGAVVVKYNFEQMLALHIRDPYE-ADSCGLCGIWDLDKSNEGPDTQYVD   450
Maristella_chicoi_se   LPGAVVVKYNFAQMLALHIRDPYE-ADSCGLCGIWDLDKSNDGPDTQYVD   450
Maristella_sp_SVD_se   LPGAVVVKYNFAQMLALHIRDPYE-ADSCGLCGIWDLDKSNDGPDTQYVD   447
Maristella_sp_SVU_se   LPGAVVVKYNFEQMLALHIRDPYE-ADSCGLCGIWDLDKSNDGPDAQHAD   450
Maristella_sp_AG_DN3   LPGAVVVKYNFAQMLALHIRDPYE-ADSCGLCGIWDLDKSNDGPDTQYAD   486
Maristella_sp_VAD_DN   LPGAVVVKYNFAQMLALHIRDPYE-ADSCGLCGIWDLDKSNDGPDTQYVD   467
Photeros_sp_WLU_DN21   LPGAVVVKYNFDQMLALHIRDPYEHGRSCGLCGLWDLDNTNDEPQRQYSE   470
Photeros_macelroyi_D   LPGAVVVKYNFDQMLALHIRDPYEHGRSCGLCGLWDLDNTNDEPQRQYSE   470
Photeros_morini_sequ   LPGAVVVKYNFDQMLALHIRDPYENGRACGLCGLWDLDNTNDEPQRQYSE   452
Photeros_annecohenae   LPGAVVVKYNFDQMLALHIRDPYEHGRSCGLCGLWDLDNTNDEPQRQYSE   470

Vargula_hilgendorfii   CALTPNPPGCTEEQKPEAERLCNNLF--DSSIDEKCNVCYKPDRIARCMY   519
Cypridina_noctiluca_   CDLTPNPPGCTEEQRPEAERLCNSLFVGQSDLDQKCNVCYKPDRVERCMY   517
Kornickeria_hastings   CEPTPNPATCTADQEAEARELCQNMF--PASLDDECDICYKSDRVERCMY   498
Vargula_tsujii_seque   CEPTPNPPTCTADKEAEARELCQNMF--PASLDDQCNICYKADRVERCMY   498
Maristella_chicoi_se   CEPTPNPPTCTADKEAEARELCQNMF--PASIDDECDICYKADRVERCMY   498
Maristella_sp_SVD_se   CEPTPNPPTCTADKEAEARELCQNMF--PASIDDECDICYKADRVERCMY   495
Maristella_sp_SVU_se   CEPTPNPPTCTADKEAEARELCQNMF--PASIDDKCNICYKADRVERCMY   498
Maristella_sp_AG_DN3   CEPTPNPPTCTADKEAEARELCQNMF--PASIDDKCNICYKADRVERCMY   534
Maristella_sp_VAD_DN   CEPTPNPPTCTADKEAEARELCQNMF--PASIDDECDICYKADRVERCMY   515
Photeros_sp_WLU_DN21   CDPTPNPPHCSAEKEAEARDLCANMF--PANLDDVCNICYNADRMKRCMY   518
Photeros_macelroyi_D   CDPTPNPPHCSAEKEAEARDLCANMF--PANLDDVCNICYNADRMKRCMY   518
Photeros_morini_sequ   CDPTPSPPHCSAEKEAEARDLCANMF--PANLDDVCNICYNADRMKRCMY   500
Photeros_annecohenae   CDPTPNPPHCSAEKEAEARDLCANMF--PANLDDVCNICYNADRMKRCMY   518

Vargula_hilgendorfii   EYCLRGQQGFCDHAWEFKKECYIKHGDTLEVPPECQ-   555
Cypridina_noctiluca_   EYCLRGQQGFCDHAWEFKKECYIKHGDTLEVPDECK-   553
Kornickeria_hastings   EYCLGGMDYFCKHAGTVIDECFVRHGDDLQLPPQCTK   535
Vargula_tsujii_seque   EYCLGGLEGFCQHAGTVIDECFVRHGDDLQYPPQCK-   534
Maristella_chicoi_se   EYCLGGLEGFRQHAGTVIDECFVRHGDDLQYPPQCK-   534
Maristella_sp_SVD_se   EYCLGGLEGFCQHAGTVIDECFVRHGDDLQYPPQCK-   531
Maristella_sp_SVU_se   EYCLGGLEAFCQHAGTVIDECFVRHGDDLQYPPQCK-   534
Maristella_sp_AG_DN3   EYCLGGLEAFCQHAGTVIDECFVRHGDDLQYPPQCK-   570
Maristella_sp_VAD_DN   EYCLGGLEGFCQHAGTVIDECFVRHGDDLQYPPQCK-   551
Photeros_sp_WLU_DN21   EYCLNGKEGICQHATSILDECFVRHGDDYKVPAICQ-   554
Photeros_macelroyi_D   EYCLNGMEGICQHANSILDECFVRHGDDYKVPAICQ-   554
Photeros_morini_sequ   EYCLNGMEGICQHANSILDECFVRHGDDYKVPAICQ-   536
Photeros_annecohenae   EYCLNGMEGICQHANSILDECFVRHGDDYKVPAICQ-   554
```

**Table S1** - Previously published emission spectra for Cypridinidae

| **Species** | **λmax(nm)** | **FWHM (nm)** | **Method** | **Light Units** | **Citation** |
|---|---|---|---|---|---|
| *Vargula hilgendorfii* | 459 | | Absolute max | Relative Quanta | (4) |
| *Vargula hilgendorfii* | 465 | 84 | Savitzky-Golay with 2°polynomial and 25 channel smoothing, slit = 1, .1 mm | Energy | (5) |
| *Vargula tsujii* | 466 | 87 | Savitzky-Golay with 2°polynomial and 25 channel smoothing, slit=1 mm | Energy | (5) |
| *Cypridina noctiluca* | 465 | | Not described | Not described | (6) |
| *Cypridina noctiluca* | 454 | | Absolute Max following FFT/LPT (>0.05) smoothing in OriginPro. N=2 | Not described | (7) |
| *Photeros shulmanae* | 473 | 80 | Cited (Widder et al. 1983) | | (8) |
| *Photeros graminicola* | 473 | 80 | Cited (Widder et al. 1983) | | (8) |

**Table S2** - Statistical results for luciferase expression *in vitro* from key comparisons

| Culture | Comparison | Test | Value | Std. Error | DF | T-value | P-value | Correted P-value (if needed) |
|---|---|---|---|---|---|---|---|---|
| Mammalian Cells | HEK & P_mor | T-test | na | na | 8.39 | -4.76 | 0.00 | 0.00 |
| Mammalian Cells | HEK & M_SVU | T-test | na | na | 11.92 | -14.91 | 0.00 | 0.00 |
| Mammalian Cells | HEK & V_tsu | T-test | na | na | 11.25 | -15.92 | 0.00 | 0.00 |
| Yeast Cells | Before & After substrate (C_noc) | T-test | na | na | 8.75 | 5.79 | 0.00 | 0.00 |
| Yeast Cells | Before & After substrate (K_has) | T-test | na | na | 4.68 | 3.18 | 0.03 | 0.11 |
| Yeast Cells | Before & After substrate (M_SVU) | T-test | na | na | 11.50 | 2.62 | 0.02 | 0.09 |
| Yeast Cells | Before & After substrate (V_tsu) | T-test | na | na | 4.60 | 4.88 | 0.01 | 0.02 |
| | **Model effects** | | | | | | | |
| Yeast Cells | Pichia after substrate addition | Linear Mixed Effect Model | 2.13 | 0.85 | 53.00 | 2.50 | 0.02 | |
| Yeast Cells | C_noc | Linear Mixed Effect Model | 3.17 | 0.98 | 19.00 | 3.23 | 0.00 | |
| Yeast Cells | K_has | Linear Mixed Effect Model | 0.88 | 1.04 | 19.00 | 0.85 | 0.41 | |
| Yeast Cells | M_SVU | Linear Mixed Effect Model | 0.57 | 0.95 | 19.00 | 0.60 | 0.55 | |
| Yeast Cells | V_tsu | Linear Mixed Effect Model | 0.49 | 1.04 | 19.00 | 0.47 | 0.65 | |
| Yeast Cells | Before substrate addition | Linear Mixed Effect Model | -0.09 | 0.28 | 40.00 | -0.31 | 0.75 | |
| Yeast Cells | C_noc * Before substrate addition | Linear Mixed Effect Model | -2.83 | 0.36 | 40.00 | -7.87 | 0.00 | |
| Yeast Cells | K_has * Before substrate addition | Linear Mixed Effect Model | -2.00 | 0.39 | 40.00 | -5.09 | 0.00 | |
| Yeast Cells | M_SVU * Before substrate addition | Linear Mixed Effect Model | -0.70 | 0.32 | 40.00 | -2.19 | 0.03 | |
| Yeast Cells | V_tsu * Before substrate addition | Linear Mixed Effect Model | -1.26 | 0.39 | 40.00 | -3.21 | 0.00 | |

**Table S3** - Means of parameters of emission spectra

| Species abbreviation | N | Lmax_Mean | Lmax_SD | FWHM_Mean | FWHM_SD |
|---|---|---|---|---|---|
| P_EGD | 4 | 468 | 0.71 | 82.05 | 1 |
| P_SFM | 4 | 467.5 | 0.71 | 82.19 | 0.28 |
| M_CON | 3 | 461.5 | 0.32 | 81.66 | 2.7 |
| M_LLL | 5 | 461 | 0.6 | 79.82 | 0.87 |
| M_MFU | 5 | 461.6 | 0.46 | 80.48 | 0.25 |
| P_ann | 3 | 468.5 | 1.8 | 85.44 | 0.32 |
| P_mor | 3 | 466.3 | 0.63 | 81.56 | 0.85 |
| M_SMU | 5 | 460.9 | 0.49 | 79.71 | 0.46 |
| K_has | 1 | 460.5 | NA | 77.98 | NA |
| M_chi | 4 | 461.3 | 0.55 | 79.24 | 0.7 |
| M_SVU | 10 | 460.9 | 0.85 | 79.43 | 1.2 |
| P_GPH | 6 | 465.6 | 0.78 | 76.77 | 0.74 |
| P_WLU | 7 | 465.9 | 1.5 | 79.08 | 1.5 |
| M_LSD | 1 | 459.4 | NA | 80.19 | NA |
| M_IR | 5 | 460.7 | 1.5 | 76.16 | 1.3 |
| M_RD | 2 | 461.5 | 2.7 | 76.54 | 1.2 |
| M_DU | 5 | 459 | 1.1 | 75.05 | 0.91 |
| V_tsu | 8 | 460.6 | 1.2 | 74.28 | 0.65 |
| K_SPU | 3 | 460.3 | 1.8 | 77.79 | 0.85 |
| V_hil | 6 | 458.7 | 2 | 78.28 | 1.8 |
| P_gra_p | 1 | 471.1 | NA | 78.58 | NA |
| V_hil_p | 1 | 460.5 | NA | 83.11 | NA |
| C_noc_p | 1 | 454.3 | NA | 79.17 | NA |

**Table S4** - All parameters measured from individual emission spectra in this study, including those removed for further analysis due to low signal:noise

| Abbre-viation | locality | genus | species | replicate | sex | preservation | source | sgMax | sgfwhm | error |
|---|---|---|---|---|---|---|---|---|---|---|
| P_EGD | Panama | Photeros | Photeros_EGD | EGD1 | male | dried | ucsb | 467.73 | 81.50 | 0.00 |
| P_EGD | Panama | Photeros | Photeros_EGD | EGD3 | male | dried | ucsb | 467.18 | 82.60 | 0.00 |
| P_EGD | Panama | Photeros | Photeros_EGD | EGD4 | male | dried | ucsb | 468.83 | 80.95 | 0.00 |
| P_EGD | Panama | Photeros | Photeros_EGD | EGD5 | male | dried | ucsb | 468.28 | 83.16 | 0.00 |
| P_SFM | Panama | Photeros | Photeros_SFM | SFM1 | male | dried | ucsb | 469.38 | 75.99 | 0.05 |
| P_SFM | Panama | Photeros | Photeros_SFM | SFM2 | male | dried | ucsb | 467.18 | 82.05 | 0.00 |
| P_SFM | Panama | Photeros | Photeros_SFM | SFM3 | male | dried | ucsb | 466.63 | 82.05 | 0.00 |
| P_SFM | Panama | Photeros | Photeros_SFM | SFM4 | male | dried | ucsb | 467.73 | 82.60 | 0.00 |
| P_SFM | Panama | Photeros | Photeros_SFM | SFM5 | male | dried | ucsb | 468.28 | 82.05 | 0.00 |
| M_CON | Panama | Contragula | contragula | cont1 | male | dried | ucsb | 461.13 | 79.82 | 0.00 |
| M_CON | Panama | Contragula | contragula | cont2 | male | dried | ucsb | 461.68 | 80.37 | 0.01 |
| M_CON | Panama | Contragula | contragula | cont3 | male | dried | ucsb | 461.68 | 84.78 | 0.00 |
| M_LLL | Panama | Maristella | Maristella_LLL | LLL1 | male | dried | ucsb | 461.13 | 79.82 | 0.00 |
| M_LLL | Panama | Maristella | Maristella_LLL | LLL2 | male | dried | ucsb | 461.68 | 80.37 | 0.00 |
| M_LLL | Panama | Maristella | Maristella_LLL | LLL3 | male | dried | ucsb | 461.13 | 78.72 | 0.00 |
| M_LLL | Panama | Maristella | Maristella_LLL | LLL4 | male | dried | ucsb | 460.03 | 79.27 | 0.00 |
| M_LLL | Panama | Maristella | Maristella_LLL | LLL5 | male | dried | ucsb | 461.13 | 80.92 | 0.00 |
| M_MFU | Panama | Maristella | Maristella_MFU | MFU1 | male | dried | ucsb | 461.13 | 80.37 | 0.00 |
| M_MFU | Panama | Maristella | Maristella_MFU | MFU2 | male | dried | ucsb | 461.68 | 80.93 | 0.00 |
| M_MFU | Panama | Maristella | Maristella_MFU | MFU3 | male | dried | ucsb | 461.13 | 80.37 | 0.00 |
| M_MFU | Panama | Maristella | Maristella_MFU | MFU4 | male | dried | ucsb | 461.68 | 80.38 | 0.00 |
| M_MFU | Panama | Maristella | Maristella_MFU | MFU5 | male | dried | ucsb | 462.23 | 80.38 | 0.00 |
| P_ann | Belize | Photeros | Photeros_annecohenae | Pann1 | unknown | live | ucsb | 466.51 | 85.25 | 0.00 |
| P_ann | Belize | Photeros | Photeros_annecohenae | Pann2 | unknown | live | ucsb | 469.81 | 85.81 | 0.01 |
| P_ann | Belize | Photeros | Photeros_annecohenae | Pann3 | unknown | live | ucsb | 469.26 | 85.25 | 0.00 |
| P_mor | Belize | Photeros | Photeros_morini | Pmor1 | male | live | ucsb | 464.32 | 79.72 | 0.05 |
| P_mor | Belize | Photeros | Photeros_morini | Pmor2 | male | live | ucsb | 467.06 | 81.37 | 0.00 |
| P_mor | Belize | Photeros | Photeros_morini | Pmor3 | male | live | ucsb | 467.61 | 83.58 | 0.04 |
| P_mor | Belize | Photeros | Photeros_morini | Pmor4 | male | live | ucsb | 465.96 | 82.49 | 0.00 |
| P_mor | Belize | Photeros | Photeros_morini | Pmor5 | male | live | ucsb | 465.96 | 80.82 | 0.00 |
| M_SMU | Panama | Maristella | Maristella_SMU | SMU1 | male | dried | ucsb | 461.13 | 79.82 | 0.00 |
| M_SMU | Panama | Maristella | Maristella_SMU | SMU2 | male | dried | ucsb | 460.58 | 79.27 | 0.00 |
| M_SMU | Panama | Maristella | Maristella_SMU | SMU3 | male | dried | ucsb | 460.58 | 79.82 | 0.00 |

| | | | | | | | | | | |
|---|---|---|---|---|---|---|---|---|---|---|
| M_SMU | Panama | Maristella | Maristella_SMU | SMU4 | male | dried | ucsb | 461.68 | 80.37 | 0.00 |
| M_SMU | Panama | Maristella | Maristella_SMU | SMU5 | male | dried | ucsb | 460.58 | 79.27 | 0.00 |
| K_has | Belize | Kornickeria | Kornickeria_hastingsi | Khas1 | male | dried | ucsb | 464.32 | 80.74 | 0.11 |
| K_has | Belize | Kornickeria | Kornickeria_hastingsi | Khas2 | male | live | ucsb | 471.46 | 75.78 | 0.09 |
| K_has | Belize | Kornickeria | Kornickeria_hastingsi | Khas3 | male | live | ucsb | 460.48 | 77.98 | 0.00 |
| M_chi | Belize | Maristella | Maristella_chicoi | MSH1 | male | live | ucsb | 461.57 | 79.10 | 0.00 |
| M_chi | Belize | Maristella | Maristella_chicoi | MSH2 | male | live | ucsb | 461.57 | 80.21 | 0.00 |
| M_chi | Belize | Maristella | Maristella_chicoi | MSH3 | male | live | ucsb | 458.83 | 79.10 | 0.02 |
| M_chi | Belize | Maristella | Maristella_chicoi | MSH4 | male | live | ucsb | 460.48 | 79.10 | 0.00 |
| M_chi | Belize | Maristella | Maristella_chicoi | MSH5 | male | live | ucsb | 461.57 | 78.55 | 0.00 |
| M_SVU | Belize | Maristella | Maristella_SVD | SVD1 | male | live | ucsb | 461.02 | 79.66 | 0.00 |
| M_SVU | Belize | Maristella | Maristella_SVD | SVD2 | male | live | ucsb | 462.12 | 82.42 | 0.00 |
| M_SVU | Belize | Maristella | Maristella_SVD | SVD3 | male | live | ucsb | 459.93 | 79.10 | 0.00 |
| M_SVU | Belize | Maristella | Maristella_SVD | SVD4 | male | live | ucsb | 461.57 | 79.65 | 0.00 |
| M_SVU | Belize | Maristella | Maristella_SVD | SVD5 | male | live | ucsb | 461.02 | 78.55 | 0.00 |
| M_SVU | Belize | Maristella | Maristella_SVU | SVU1 | male | live | ucsb | 461.02 | 79.10 | 0.00 |
| M_SVU | Belize | Maristella | Maristella_SVU | SVU2 | male | live | ucsb | 462.12 | 79.66 | 0.00 |
| M_SVU | Belize | Maristella | Maristella_SVU | SVU3 | male | live | ucsb | 459.93 | 77.99 | 0.00 |
| M_SVU | Belize | Maristella | Maristella_SVU | SVU4 | male | live | ucsb | 460.48 | 79.10 | 0.00 |
| M_SVU | Belize | Maristella | Maristella_SVU | SVU5 | male | live | ucsb | 459.93 | 79.10 | 0.00 |
| P_GPH | Roatan | Photeros | Photeros_GPH | GPH1 | male | live | ucsb | 466.19 | 75.75 | 0.01 |
| P_GPH | Roatan | Photeros | Photeros_GPH | GPH2 | male | live | ucsb | 466.19 | 75.75 | 0.08 |
| P_GPH | Roatan | Photeros | Photeros_GPH | GPH3 | male | live | ucsb | 465.63 | 76.30 | 0.01 |
| P_GPH | Roatan | Photeros | Photeros_GPH | GPH4 | male | live | ucsb | 467.84 | 75.21 | 0.06 |
| P_GPH | Roatan | Photeros | Photeros_GPH | GPH5 | male | live | ucsb | 466.74 | 77.42 | 0.00 |
| P_GPH | Roatan | Photeros | Photeros_GPH | GPH6 | male | live | ucsb | 465.63 | 77.41 | 0.00 |
| P_GPH | Roatan | Photeros | Photeros_GPH | GPH7 | male | live | ucsb | 464.53 | 76.31 | 0.00 |
| P_GPH | Roatan | Photeros | Photeros_GPH | GPH8 | male | live | ucsb | 465.08 | 77.42 | 0.01 |
| P_WLU | Roatan | Photeros | Photeros_WLU | WLU1 | male | live | ucsb | 466.19 | 79.08 | 0.00 |
| P_WLU | Roatan | Photeros | Photeros_WLU | WLU2 | male | live | ucsb | 463.98 | 77.41 | 0.00 |
| P_WLU | Roatan | Photeros | Photeros_WLU | WLU3 | male | live | ucsb | 466.19 | 78.52 | 0.00 |
| P_WLU | Roatan | Photeros | Photeros_WLU | WLU4 | male | live | ucsb | 464.53 | 78.52 | 0.00 |
| P_WLU | Roatan | Photeros | Photeros_WLU | WLU5 | male | live | ucsb | 466.74 | 80.19 | 0.00 |
| P_WLU | Roatan | Photeros | Photeros_WLU | WLU6 | male | live | ucsb | 465.63 | 77.97 | 0.00 |
| P_WLU | Roatan | Photeros | Photeros_WLU | WLU7 | male | dried | ucsb | 466.74 | 78.53 | 0.03 |
| P_WLU | Roatan | Photeros | Photeros_WLU | WLU8 | male | dried | ucsb | 468.40 | 81.87 | 0.01 |
| M_LSD | PR_USA | Maristella | Maristella_LSD | LSD1 | male | live | ucsb | 459.38 | 80.19 | 0.02 |
| M_LSD | PR_USA | Maristella | Maristella_LSD | LSD2 | male | live | ucsb | 459.93 | 79.08 | 0.03 |
| M_IR | Roatan | Maristella | Maristella_IR | IR1 | male | dead | ucsb | 459.56 | 75.71 | 0.00 |

| | | | | | | | | | | |
|---|---|---|---|---|---|---|---|---|---|---|
| M_IR | Roatan | Maristella | Maristella_IR | IR2 | male | dead | ucsb | 459.56 | 76.82 | 0.00 |
| M_IR | Roatan | Maristella | Maristella_IR | IR3 | male | live | ucsb | 466.74 | 83.46 | 0.13 |
| M_IR | Roatan | Maristella | Maristella_IR | IR4 | male | dead | ucsb | 459.56 | 74.60 | 0.01 |
| M_IR | Roatan | Maristella | Maristella_IR | IR5 | male | dead | ucsb | 462.32 | 77.93 | 0.01 |
| M_IR | Roatan | Maristella | Maristella_IR | IR6 | male | dead | ucsb | 462.32 | 75.71 | 0.01 |
| M_RD | Roatan | Maristella | Maristella_RD | RD1 | male | dead | ucsb | 463.42 | 75.71 | 0.02 |
| M_RD | Roatan | Maristella | Maristella_RD | RD2 | male | dead | ucsb | 459.56 | 77.38 | 0.01 |
| M_RD | Roatan | Maristella | Maristella_RD | RD3 | male | dead | ucsb | 466.74 | 77.92 | 0.10 |
| M_RD | Roatan | Maristella | Maristella_RD | RD4 | male | dead | ucsb | 465.08 | 78.48 | 0.03 |
| M_RD | Roatan | Maristella | Maristella_RD | RD5 | male | dead | ucsb | 465.63 | 75.16 | 0.04 |
| M_DU | Roatan | Maristella | Maristella_DU | DU1 | male | live | ucsb | 460.11 | 75.16 | 0.00 |
| M_DU | Roatan | Maristella | Maristella_DU | DU2 | male | live | ucsb | 459.56 | 75.16 | 0.00 |
| M_DU | Roatan | Maristella | Maristella_DU | DU3 | male | live | ucsb | 457.35 | 75.71 | 0.00 |
| M_DU | Roatan | Maristella | Maristella_DU | DU4 | male | live | ucsb | 458.45 | 73.49 | 0.01 |
| M_DU | Roatan | Maristella | Maristella_DU | DU5 | male | live | ucsb | 459.56 | 75.71 | 0.00 |
| V_tsu | CA_USA | Fred | Vargula_tsujii | Vtsu1 | female | live | ucsb | 462.12 | 74.14 | 0.01 |
| V_tsu | CA_USA | Fred | Vargula_tsujii | Vtsu2 | female | live | ucsb | 461.02 | 74.70 | 0.00 |
| V_tsu | CA_USA | Fred | Vargula_tsujii | Vtsu3 | female | live | ucsb | 462.12 | 74.14 | 0.01 |
| V_tsu | CA_USA | Fred | Vargula_tsujii | Vtsu4 | male | live | ucsb | 458.83 | 74.14 | 0.00 |
| V_tsu | CA_USA | Fred | Vargula_tsujii | Vtsu5 | male | live | ucsb | 459.93 | 74.14 | 0.00 |
| V_tsu | CA_USA | Fred | Vargula_tsujii | Vtsu6 | male | live | ucsb | 459.93 | 73.03 | 0.01 |
| V_tsu | CA_USA | Fred | Vargula_tsujii | Vtsu7 | male | live | ucsb | 459.93 | 75.25 | 0.00 |
| V_tsu | CA_USA | Fred | Vargula_tsujii | Vtsu8 | male | live | ucsb | 461.02 | 74.70 | 0.01 |
| K_WCU | PR_USA | Kornickeria | Kornickeria_WCU | WCU1 | male | live | ucsb | 462.12 | 81.78 | 0.14 |
| K_WCU | PR_USA | Kornickeria | Kornickeria_WCU | WCU2 | male | live | ucsb | 458.83 | 79.07 | 0.03 |
| K_WCU | PR_USA | Kornickeria | Kornickeria_WCU | WCU3 | male | live | ucsb | 463.22 | 82.36 | 0.08 |
| K_WCU | PR_USA | Kornickeria | Kornickeria_WCU | WCU5 | male | live | ucsb | 450.06 | 74.09 | 0.08 |
| K_SPU | PR_USA | Kornickeria | Kornickeria_SPU | SPU1 | male | live | ucsb | 457.19 | 76.89 | 0.02 |
| K_SPU | PR_USA | Kornickeria | Kornickeria_SPU | SPU2 | male | live | ucsb | 458.28 | 76.87 | 0.00 |
| K_SPU | PR_USA | Kornickeria | Kornickeria_SPU | SPU3 | male | live | ucsb | 461.57 | 78.54 | 0.01 |
| K_SPU | PR_USA | Kornickeria | Kornickeria_SPU | SPU4 | male | live | ucsb | 454.99 | 73.56 | 0.07 |
| K_SPU | PR_USA | Kornickeria | Kornickeria_SPU | SPU5 | male | live | ucsb | 460.48 | 81.85 | 0.08 |
| K_SPU | PR_USA | Kornickeria | Kornickeria_SPU | SPU6 | male | live | ucsb | 461.02 | 77.97 | 0.01 |
| K_SPU | PR_USA | Kornickeria | Kornickeria_SPU | SPU7 | male | live | ucsb | 451.16 | 71.31 | 0.13 |
| V_hil | Japan | Vargula | Vargula_hilgendorfii | Vhil012317 1 | unknown | dried | ucsb | 460.48 | 76.88 | 0.01 |
| V_hil | Japan | Vargula | Vargula_hilgendorfii | Vhil012317 2 | unknown | dried | ucsb | 459.38 | 77.98 | 0.00 |
| V_hil | Japan | Vargula | Vargula_hilgendorfii | Vhil012317 3 | unknown | dried | ucsb | 459.38 | 74.68 | 0.04 |

| | | | | | | | | | | |
|---|---|---|---|---|---|---|---|---|---|---|
| V_hil | Japan | Vargula | Vargula_hilgendorfii | Vhil0907161 | unknown | dried | ucsb | 459.66 | 80.87 | 0.03 |
| V_hil | Japan | Vargula | Vargula_hilgendorfii | Vhil0907162 | unknown | dried | ucsb | 460.21 | 77.54 | 0.06 |
| V_hil | Japan | Vargula | Vargula_hilgendorfii | Vhil0907163 | unknown | dried | ucsb | 460.21 | 80.87 | 0.03 |
| V_hil | Japan | Vargula | Vargula_hilgendorfii | Vhil0907164 | unknown | dried | ucsb | 456.33 | 78.10 | 0.02 |
| V_hil | Japan | Vargula | Vargula_hilgendorfii | Vhil0907165 | unknown | dried | ucsb | 456.33 | 76.99 | 0.01 |
| V_hil | Japan | Vargula | Vargula_hilgendorfii | Vhil0907166 | unknown | dried | ucsb | 460.76 | 79.76 | 0.04 |
| V_hil | Japan | Vargula | Vargula_hilgendorfii | Vhil100520161 | unknown | dried | ucsb | 458.28 | 81.85 | 0.01 |
| V_hil | Japan | Vargula | Vargula_hilgendorfii | Vhil0920161 | unknown | dried | ucsb | 465.19 | 73.11 | 0.02 |
| V_hil | Japan | Vargula | Vargula_hilgendorfii | Vhil0920162 | unknown | dried | ucsb | 465.19 | 72.01 | 0.02 |
| V_hil | Japan | Vargula | Vargula_hilgendorfii | Vhil0920163 | unknown | dried | ucsb | 461.32 | 72.01 | 0.03 |
| V_hil | Japan | Vargula | Vargula_hilgendorfii | Vhil_Japan | unknown | live | Japan | 461.14 | 77.87 | 0.00 |
| P_gra_p | Published | Photeros | Photeros_gramminicola | Pgra_huvard | unknown | unknown | published | 471.14 | 78.58 | 0.32 |
| V_hil_p | Published | Vargula | Vargula_hilgendorfii | Vhil_tsuji | unknown | unknown | published | 460.54 | 83.11 | 0.33 |
| C_noc_p | Published | Cypridina | Cypridina_noctiluca | Cnoc_ohmiya | unknown | unknown | published | 454.27 | 79.17 | 0.06 |

**Table S5** - Collection localities, size measurements, and accession numbers for vouchers for specimens used in this study

| **Inferred genus** | **Species or field code** | **Locality** | **Latitude** | **Longitude** | **Carapace length** | **Carapace height** | **Carapace ratio** | **BioProject** | **Luciferase Genbank Accession** | **SRA Accession** | **RNA Isolation** | **Library Preparation** | **Sequencing Instrument** |
|---|---|---|---|---|---|---|---|---|---|---|---|---|---|
| Photeros | EGD | Bocas del Toro, Panama | 9.331385 | -82.25434 | 1.625 | 1.044 | 1.56 | | N/A | N/A | | | |
| Photeros | SFM | Bocas del Toro, Panama | 9.331326 | -82.25327 | 1.703 | 1.077 | 1.58 | | N/A | N/A | | | |
| Photeros | annecohenae | Southwater Caye, Belize | 16.8116 | -88.08243 | 1.623 | 1.02 | 1.59 | PRJNA589015 | | SRR10860880 | Qiagen Rneasy Fibrous Tissue Mini Kit | Illumina TruSeq v3 | Illumina HiSeq 2500 |
| Photeros | morini | Southwater Caye, Belize | 16.8116 | -88.08243 | 2.056 | 1.282 | 1.6 | PRJNA589015 | | SRR10860877 | Trizol | NEB Ultra RNA Library Prep Kit for Illumina | Illumina HiSeq 1500 |
| Photeros | morini | Southwater Caye, Belize | 16.8116 | -88.08243 | 2.056 | 1.282 | 1.6 | PRJNA589015 | | SRR10860876 | Qiagen Rneasy | Illumina TruSeq v2 | Illumina HiSeq 1500 |
| Photeros | GPH | Roatan, Honduras | 16.40272 | -86.409 | 1.683 | 1.038 | 1.62 | | N/A | N/A | | | |
| Photeros | WLU | Roatan, Honduras | 16.35806 | -86.43291 | 1.978 | 1.222 | 1.62 | PRJNA589015 | | SRR10811635 | Trizol | NEBNextUltra IIRNA Library Prep Kit for Illumina | NextSeq 500 |
| Photeros | mcelroyi | Discovery Bay, Jamaica | | | | | | PRJNA589015 | | SRR10811638 | Qiagen RNA | Illumina TruSeq v3 | Illumina HiSeq 1500 |
| Photeros | mcelroyi | Discovery Bay, Jamaica | | | | | | PRJNA589015 | | SRR10811637 | Qiagen RNA | Illumina TruSeq v3 | Illumina HiSeq 1500 |
| Photeros | mcelroyi | Discovery Bay, Jamaica | | | | | | PRJNA589015 | | SRR10811645 | Trizol | NEBNext Ultra RNA Library Prep Kit for Illumina | Illumina HiSeq 1500 |

| | | | | | | | | | | | | | |
|---|---|---|---|---|---|---|---|---|---|---|---|---|---|
| Maristella | MFU | Bocas del Toro, Panama | 9.331326 | -82.253267 | 1.657 | 1.002 | 1.65 | | N/A | N/A | | | |
| Maristella | IR | Roatan, Honduras | 16.35806 | -86.43291 | 2.282 | 1.386 | 1.65 | | N/A | N/A | | | |
| Maristella | SVU (MWU) | Southwater Caye, Belize | 16.8116 | -88.08243 | 2.172 | 1.305 | 1.66 | PRJNA589015 | | SRR10811644 | Trizol | NEBNext Ultra RNA Library Prep Kit for Illumina | Illumina HiSeq 1500 |
| Maristella | SVU (MWU) | Southwater Caye, Belize | 16.8116 | -88.08243 | 2.172 | 1.305 | 1.66 | PRJNA589015 | | SRR10811643 | Qiagen RNA | Illumina TruSeq v2 | Illumina HiSeq 1000 |
| Maristella | SVD | Southwater Caye, Belize | 16.8116 | -88.08243 | | | | PRJNA589015 | | SRR10860875 | Trizol | NEB Ultra RNA Library Prep Kit for Illumina | Illumina HiSeq 1500 |
| Maristella | SVD | Southwater Caye, Belize | 16.8116 | -88.08243 | | | | PRJNA589015 | | SRR10860874 | Qiagen Rneasy | Illumina TruSeq v2 | Illumina HiSeq 1500 |
| Maristella | DU | Roatan, Honduras | 16.35806 | -86.43291 | 1.631 | 0.981 | 1.66 | | N/A | N/A | | | |
| Maristella | RD | Roatan, Honduras | 16.358061 | -86.432906 | 1.637 | 0.98 | 1.67 | | N/A | N/A | | | |
| Maristella | LLL | Bocas del Toro, Panama | 9.331707 | -82.255633 | 1.775 | 1.057 | 1.68 | | N/A | N/A | | | |
| Maristella | SMU | Bocas del Toro, Panama | 9.331707 | -82.255633 | 2.162 | 1.289 | 1.68 | | N/A | N/A | | | |
| Maristella | chicoi | Southwater Caye, Belize | 16.8116 | -88.08243 | 1.624 | 0.963 | 1.69 | PRJNA589015 | N/A | SRR10811640 | Trizol | NEBNext Ultra RNA Library Prep Kit for Illumina | Illumina HiSeq 1500 |
| Maristella | chicoi | Southwater Caye, Belize | 16.8116 | -88.08243 | 1.624 | 0.963 | 1.69 | PRJNA589015 | N/A | SRR10811639 | Qiagen RNA | Illumina TruSeq v2 | Illumina HiSeq 1500 |
| Maristella | VAD | Discovery Bay, Jamaica | | | | | | PRJNA589015 | | SRR10811642 | Qiagen RNA | Illumina TruSeq v3 | Illumina HiSeq 1500 |
| Maristella | VAD | Discovery Bay, Jamaica | | | | | | PRJNA589015 | | SRR10811641 | Trizol | NEBNextUltraRNA Library Prep Kit | Illumina MiSeq |

| | | | | | | | | | | | | | |
|---|---|---|---|---|---|---|---|---|---|---|---|---|---|
| | | | | | | | | | | | | for Illumin | |
| Maristella | AG | Roatan, Honduras | 16.358061 | -86.432906 | | | | PRJNA589015 | | SRR10811636 | Trizol | NEBNextUltra IIRNA Library Prep Kit for Illumina | NextSeq 500 |
| C-group | CONT | Bocas del Toro, Panama | 9.331707 | -82.255633 | 1.846 | 1.066 | 1.73 | | N/A | N/A | | | |
| "Vargula" | tsujii | Catalina Island, CA, USA | 33.445139 | -118.4845 | 1.581 | 0.913 | 1.73 | PRJNA287212 | | SRR1269674 | | | |
| Kornickeria | SPU | Isla Magueyes, PR, USA | 17.961089 | -67.052156 | 1.383 | 0.78 | 1.77 | | N/A | N/A | | | |
| Kornickeria | hastingsi carriebowae | Southwater Caye, Belize | 16.8116 | -88.08243 | 1.799 | 1.014 | 1.77 | PRJNA589015 | | SRR10860879 | Qiagen Rneasy | Unknown; amplified by Novogene Co (Davis CA) | Illumina HiSeq 1500 |
| Kornickeria | hastingsi carriebowae | Southwater Caye, Belize | 16.8116 | -88.08243 | 1.799 | 1.014 | 1.77 | PRJNA589015 | | SRR10860878 | Qiagen Rneasy | Illumina TruSeq v2 | Illumina HiSeq 1500 |
| Maristella | LSD | Isla Magueyes, PR, USA | 17.961089 | -67.052156 | | | | | N/A | | | | |
| "Vargula" | hilgendorfii | Carolina Biological Purchase | N/A | | N/A | | | | AAA30332 | | | | |
| Cypridina | noctiluca | NCBI | | | | | | | BBG57195 | | | | |

**Table S6** - Luciferase-specific primers to amplify from cDNA

| Primer name | Target species | Sequence | Purpose |
|---|---|---|---|
| KHC Forward | *Kornickeria hastingsi carriebowae* | GGACTCGAGAAGAGAGAGGCTAAAGATTGT TTTGAATCATCTTTCC | Amplify from cDNA |
| KHC Reverse | *Kornickeria hastingsi carriebowae* | AGCGGCCGCTTTGGTGCATTGAGGTGG | Amplify from cDNA |
| PMO Forward | *Photeros morini* | TAACTCGAGAAGAGAGAGGCTCAAGAATGC GCTCAGACA | Amplify from cDNA |
| PMO Reverse | *Photeros morini* | AGCGGCCGCCTGGCATATGGCTGGTAC | Amplify from cDNA |
| SVU Forward | SVU (undescribed) | GGGCTCGAGAAGAGAGAGGCTCAAGATTGT TATGAATTCACA | Amplify from cDNA |
| SVU Reverse | SVU (undescribed) | AGCGGCCGCTTTACACTGAGGGGGATA | Amplify from cDNA |
| SVD Forward | SVD (undescribed) | GTGCTCGAGAAGAGAGAGGCTGAAGATTGT TATGAATTCACATG | Amplify from cDNA |
| SVD Reverse | SVD (undescribed) | AGCGGCCGCTTTACACTGAGGTGGATAC | Amplify from cDNA |
| MSH Forward | *Maristella chicoi* | CGGCTCGAGAAGAGAGAGGCTGAAGATTGT TATGAATTCACAT | Amplify from cDNA |
| MSH Reverse | *Maristella chicoi* | AGCGGCCGCTTTACACTGAGGGGGATAC | Amplify from cDNA |
| VTS Forward | *Vargula tsujii* | CCGCTCGAGAAGAGAGAGGCTCAAGATTGT TATGAATCAACATG | Amplify from cDNA |
| VTS Reverse | *Vargula tsujii* | AGCGGCCGCTTTACACTGAGGAGGATACT | Amplify from cDNA |
| 454-Forward-Vt | *Vargula tsujii* | CTCGAGATCAGTCGAGAAACGAAACGTGAT A | Amplify from cDNA |
| 454-Reverse-Vt | *Vargula tsujii* | GAATTCCTTTACACTGAGGGGGATACTG | Amplify from cDNA |
| AOX Forward | n/a | GACTGGTTCCAATTGACAAGC | Sequence from clones |
| AOX Reverse | n/a | GCAAATGGCATTCTGACATCC | Sequence from clones |
